# A feedback loop between heterochromatin and the nucleopore complex controls germ-cell to oocyte transition during *Drosophila* oogenesis

**DOI:** 10.1101/2021.10.31.466575

**Authors:** Kahini Sarkar, Noor M Kotb, Alex Lemus, Elliot T Martin, Alicia McCarthy, Justin Camacho, Ayman Iqbal, Alex M. Valm, Morgan A Sammons, Prashanth Rangan

## Abstract

Germ cells differentiate into oocytes that become totipotent upon fertilization. How the highly specialized oocyte acquires this distinct cell fate is poorly understood. During *Drosophila* oogenesis, H3K9me3 histone methyltransferase SETDB1 translocates from the cytoplasm to the nucleus of germ cells concurrent with oocyte specification. Here, we discovered that nuclear SETDB1 is required to silence a cohort of differentiation-promoting genes by mediating their heterochromatinization. Intriguingly, SETDB1 is also required for the upregulation of 18 of the ~30 nucleoporins (Nups) that comprise the nucleopore complex (NPC). NPCs in turn anchor SETDB1-dependent heterochromatin at the nuclear periphery to maintain H3K9me3 and gene silencing in the egg chambers. Aberrant gene expression due to loss of SETDB1 or Nups results in loss of oocyte identity, cell death and sterility. Thus, a feedback loop between heterochromatin and NPCs promotes transcriptional reprogramming at the onset of oocyte specification that is critical to establish oocyte identity.

## Introduction

Germ cells give rise to gametes that upon fertilization launch the next generation (Cinalli et al., 2008; Seydoux and Braun, 2006; Spradling et al., 2011). In the gonad, germ cells become germline stem cells (GSCs) that self-renew and differentiate to give rise to sperm or an oocyte (Gilboa and Lehmann, 2004; Kershner et al., 2013; Ko et al., 2010; Lesch and Page, 2012; Reik and Surani, 2015; Seydoux and Braun, 2006). The oocyte, upon fertilization or by parthenogenesis, can differentiate into every cell lineage in the adult organism and thus has a capacity to be totipotent (Ben-Ami and Heller, 2005; Lehmann, 2012; Riparbelli et al., 2017; Yuan and Yamashita, 2010). The gene regulatory mechanisms that enable the transition from germ cells to oocytes are not fully understood.

*Drosophila* has a well-characterized transition from germline stem cell (GSC) to an oocyte (Dansereau and Lasko, 2008; Gilboa and Lehmann, 2004; Spradling et al., 2011; Allan C Spradling, 1993). *Drosophila* ovaries comprise individual units called ovarioles that house the GSCs in a structure called the germarium (**Figure 1A-A1**) (Lehmann, 2012; Xie and Spradling, 2000). GSC division results in a new GSC (self-renewal) and a cystoblast, which differentiates via incomplete mitotic divisions, giving rise to 2-, 4-, 8- and 16-cell cysts (**Figure 1A1**) (Chen and McKearin, 2003a, 2003b; Xie, 2013). One of these 16 cells is specified as the oocyte whereas the other 15 cells become nurse cells (Huynh and St Johnston, 2004; Koch et al., 1967; Navarro et al., 2004). Somatic cells envelop the nurse cells and the specified oocyte to form an egg chamber (**Figure 1A1**) (Xie and Spradling, 2000). The nurse cells produce mRNAs, called maternal mRNAs, that are deposited into the specified oocyte mediated by an RNA binding protein, Egalitarian (Egl) (Blatt et al., 2020; Kugler and Lasko, 2009; Lilly and Spradling, 1996; Mach and Lehmann, 1997; Navarro et al., 2004; A C Spradling, 1993). An inability to specify or maintain the oocyte fate leads to death of the egg chamber mid-oogenesis and, in turn, sterility (Blatt et al., 2021; Navarro et al., 2004).

**Figure1:**
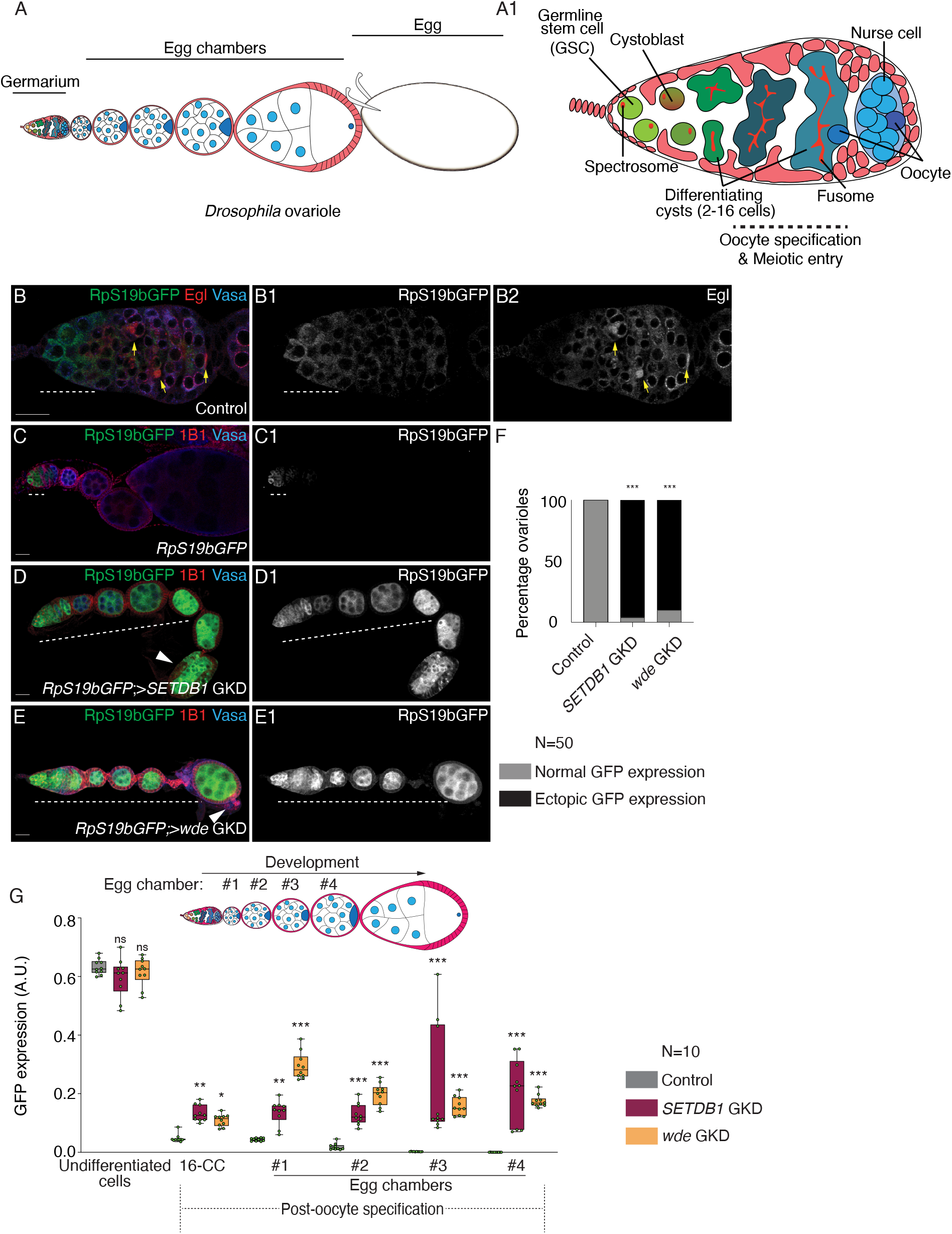
*SETDB1* and *windei* are required for silencing *RpS19b* reporter during oogenesis. (A) A schematic of a *Drosophila* ovariole. The ovariole consists of germarium and egg chambers corresponding to distinct developmental stages. Egg chambers are connected by somatic cells (light red). During development, egg chambers grow and eventually give rise to a mature egg (white). (A1) A schematic of a *Drosophila* germarium, where germline stem cells (GSCs; light green) are at close proximity to somatic niche (red). The GSC divides to give rise to daughter cells called cystoblasts (dark green) which turns on a differentiation program. Both GSCs and cystoblasts are marked by spectrosomes (red). Cystoblasts undergo four incomplete mitotic divisions, giving rise to 2-, 4-, 8-, and 16-cell cysts (green), marked by fusomes (red). During the cyst stages germ cells progress through meiotic cell cycle (prophase I). Upon 16-cell cyst formation, a single cell is specified as the oocyte (dark blue) while the other 15 cells become nurse cells (light blue). (B-B2) Confocal images of a germarium of a fly carrying *RpS19b-GFP* reporter transgene stained for GFP (green, right grayscale), Egl (red, right grayscale) and Vasa (blue). GFP is expressed in the single cells undifferentiated stages and early cyst stages (white dashed line), while Egl is expressed in the differentiated cysts and localized to the specified oocyte (yellow arrows). (C-E) Ovariole of control *RpS19b-GFP* (D-D1), GKD of *SETDB1* (E-E1) and *wde* (F-F1) stained for GFP (green, right grayscale), Vasa (blue) and 1B1 (red). Depletion of these genes resulted in characteristic phenotype of egg chambers not growing and mid oogenesis death (white solid arrows). In addition, ectopic expression of *RpS19b-*GFP was observed in the egg chambers (white dashed line). (F) Quantification of ectopic RpS19b-GFP expression upon GKD of *SETDB1* or *wde* compared to control ovaries (N= 50 ovarioles; 96% in *SETDB1* GKD and 90% in *wde* GKD compared to 0% in control.) Statistical analysis was performed with Fisher’s exact test on ectopic GFP expression; *** = p<0.001. (G) Arbitrary units (A.U.) quantification of RpS19b-GFP expression in the germarium and egg chambers during development upon GKD of *SETDB1* (magenta) or *wde* (orange) compared to control ovaries (black). GFP is expressed higher in single cells in the germarium, decreases in the cyst stages, and then attenuated upon egg chamber formation. In *SETDB1* and *wde* GKD, GFP expression persists in the egg chambers. Statistical analysis was performed with Dunnett’s multiple comparisons test; N= 10 ovarioles; ns = p>0.05, *=p<0.05, ** = p<0.01, *** = p<0.001. Scale bars are 15 micron.

The transition from GSC to an oocyte requires dynamic changes in gene expression that promote progressive differentiation (Flora et al., 2017). Once a GSC gives rise to the cystoblast, it expresses differentiation factor Bag of marbles (Bam), promoting its differentiation to an 8-cell cyst (McKearin and Ohlstein, 1995; McKearin and Spradling, 1990). In the 8-cell cyst, the expression of the RNA binding fox-1 homolog 1 (Rbfox1) is required to mediate transition into the 16-cell cyst stage, allowing for an oocyte to be specified (Carreira-Rosario et al., 2016). Translation of Rbfox1 requires increased levels of ribosomal small subunit protein 19 (RpS19) accomplished in part by expression of the germline specific paralog *RpS19b* in the undifferentiated and early differentiating stages (McCarthy et al., 2019). During differentiation, the germline also initiates meiotic recombination mediated by the synaptonemal complex consisting of proteins such as Sisters Unbound (Sunn), Corona (Cona) and Orientation Disruptor (Ord) (Ables, 2015; Cahoon and Hawley, 2016; Hughes et al., 2018; Orr-Weaver, 1995; Page and Hawley, 2001). More than one cell in the cyst stage initiates recombination but as oocyte differentiation proceeds, only the specified oocyte retains the synaptonemal complex (**Figure 1A1**) (Ables, 2015; Orr-Weaver, 1995; Page and Hawley, 2001). After oocyte-specification, the levels of mRNAs encoding *RpS19b* and some synaptonemal complex proteins are diminished, suggesting early oogenesis genes are no longer expressed (McCarthy et al., 2019). How the expression of these early oogenesis genes is attenuated is not known.

In *Drosophila*, the <u>SET</u> <u>D</u>omain <u>B</u>ifurcated Histone Lysine Methyltransferase 1 (SETDB1) (also called Eggless) is required for deposition of gene silencing Histone H3 Lysine 9 trimethylation (H3K9me3) marks and heterochromatin formation (Clough et al., 2014, 2007; Rangan et al., 2011; Yoon et al., 2008). SETDB1 is expressed throughout *Drosophila* oogenesis, but as the oocyte is specified, it shifts from a cytoplasmic to predominantly nuclear localization (Clough et al., 2007). A conserved cofactor called Windei (Wde) is required for either nuclear translocation, nuclear stability, or targeting of SETDB1 to its target loci (Koch et al., 2009; Osumi et al., 2019). Loss of *SETDB1* during germline development results in an accumulation of undifferentiated cells (Rangan et al., 2011; Smolko et al., 2018). In addition, loss of *SETDB1* and *wde* also result in egg chambers that do not grow in size and die mid-oogenesis (Clough et al., 2014; Koch et al., 2009). *SETDB1* is known to be required for silencing transposons and male-specific transcripts in the female germline (Czech et al., 2018; Rangan et al., 2011; Smolko et al., 2018). However, neither the upregulation of transposons nor male-specific genes in female germline result in egg chambers that do not grow in size (Malone et al., 2009; Shapiro-Kulnane et al., 2015; Smolko et al., 2020). Together these data suggests that SETDB1 silences a yet-unidentified group of genes to promote oogenesis.

Here, we find that genes that are expressed in early stages of oogenesis, including genes that promote oocyte differentiation and synaptonemal complex formation, are silenced upon oocyte specification, via a feedback loop between SETDB1-mediated heterochromatin and the nucleopore complex (NPC). Inability to silence these differentiation-promoting genes due to loss of either SETDB1 or members of the NPC results in loss of oocyte identity and death. Several aspects of germ cell differentiation have been studied and have been implicated in loss of fertility in sexually reproducing organisms. Our work indicates that a previously unappreciated broad transcriptional reprogramming silences critical aspects of the germ cell differentiation program at the onset of oocyte specification and is essential to promote oocyte identity.

## Results

### *SETDB1* promotes silencing of *RpS19b* reporter at the onset of oocyte specification

We hypothesized that the expression of early oogenesis mRNAs such as *RpS19b* is silenced upon oocyte specification. To monitor *RpS19b* expression, we used a reporter that expresses an RpS19b-GFP fusion from the endogenous *RpS19b* promoter. This RpS19b-GFP shows high expression in the germarium and attenuated expression post-oocyte specification and in the subsequent egg chambers, consistent with its endogenous *RpS19b* mRNA expression pattern (**Figure 1B-C1, G**) (Jevitt et al., 2020; McCarthy et al., 2019).

Using a previously characterized hemagglutinin (HA) tagged endogenous SETDB1, we found that a large fraction of SETDB1 translocates from the cytoplasm to the nucleus concurrent with oocyte specification (**Figure S1A-A3**) (Seum et al., 2007). To test if *SETDB1* is required for the silencing of *RpS19b* (Clough et al., 2014, 2007), we performed germline knockdown (GKD) of *SETDB1*, in the background of *RpS19b-GFP* reporter. We detected the germline, RpS19b-GFP, and spectrosomes/fusomes/somatic cell membrane in ovaries by immunostaining for Vasa, GFP, and 1B1, respectively (Lasko and Ashburner, 1988; Zaccai and Lipshitz, 1996). We found that, compared to the control, GKD of *SETDB1* resulted in ectopic RpS19b-GFP protein expression in the differentiated egg chambers without affecting levels in the undifferentiated stages (**Figure 1C-G; Figure S1B**). Thus, *SETDB1* is required for repression of *RpS19b-GFP* reporter in the differentiated egg chambers.

To determine if nuclear SETDB1 was required to repress *RpS19b-GFP* post-oocyte specification, we depleted *wde* in the germline and independently assayed for SETDB1 nuclear localization, H3K9me3, and RpS19b-GFP (**Figure S1C**). GKD of *wde* resulted in loss of nuclear SETDB1 in the differentiated stages of oogenesis without affecting cytoplasmic levels in the undifferentiated stages (**Figure S1D-F**). Whereas GKD of *SETDB1* reduced H3K9me3 throughout oogenesis, GKD of *wde* reduced H3K9me3 only in the differentiated egg chambers but not in the undifferentiated stages (**Figure S1G-J**). We found that GKD of *wde*, like GKD of *SETDB1*, results in ectopic RpS19b-GFP protein expression in the egg chambers without affecting levels in the undifferentiated stages (**Figure 1C-G**). In addition to upregulation of RpS19b-GFP, GKD of both *SETDB1* and *wde* resulted in egg chambers that did not grow in size and died mid-oogenesis as previously reported (**Figure S1K**) (Clough et al., 2014; Koch et al., 2009). Thus, repression of the *RpS19b-GFP* reporter in the differentiated egg chambers requires nuclear SETDB1.

### SETDB1 and Wde repress genes that are primarily expressed prior to oocyte specification

To determine if SETDB1 and Wde repress other differentiation-promoting genes in addition to *RpS19b*, we performed RNA Sequencing (RNA-seq). We compared ovaries from *SETDB1-* and *wde-* GKD flies to ovaries from wild-type (WT) flies, including young virgin flies which lack late-stage egg chambers. Principal component analysis of the RNA-seq data revealed that *SETDB1* and *wde* ovary transcriptomes closely resembles young virgin WT rather than adult WT (**Figure S2A**). Using a 1.5-fold cut off (Fold Change (FC) ≥ |1.5|) and False Discovery Rate (FDR)<0.05, we found that compared to young virgin WT control, 2316 genes were upregulated and 1972 were downregulated in *SETDB1* GKD ovaries, and 1075 genes were upregulated and 442 were downregulated in *wde-*GKD ovaries (**Figure 2A-B**) **(Supplemental Table 1)**. Moreover, comparing *wde*- to *SETDB1*-GKD ovaries showed significant overlap of the upregulated (80%) and downregulated (75%) transcripts, suggesting that *SETDB1* and *Wde* co-regulate a cohort of genes during oogenesis (**Figure 2C; Figure S2B**).

**Figure 2:**
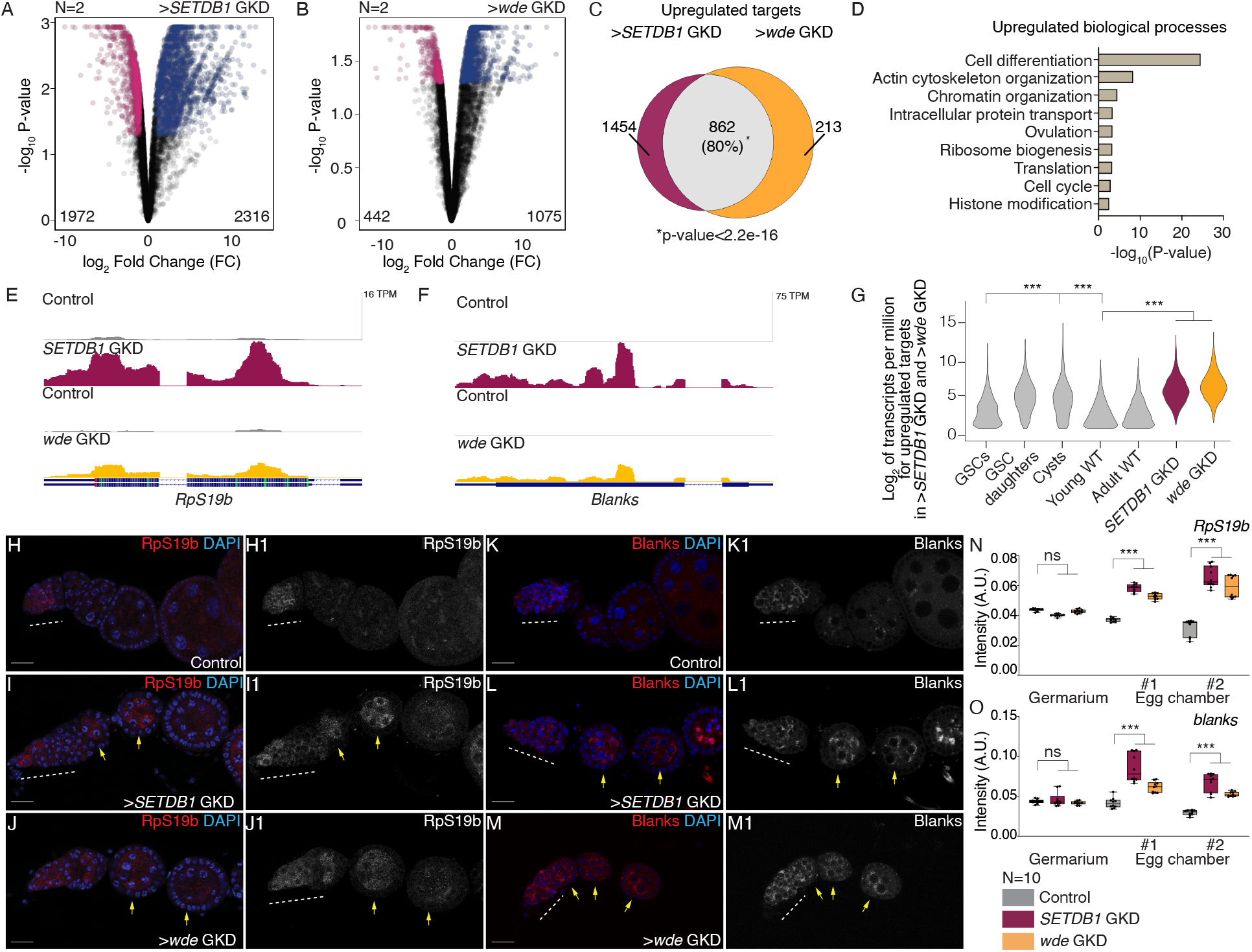
SETDB1/Wde represses a cohort of early oogenesis gene. (A-B) Volcano plots of –Log_10_P-value vs. Log2Fold Change (FC) of (A) *SETDB1* and (B) *wde* GKD ovaries compared to control *UAS-Dcr2;NG4NGT* flies. Pink dots represent significantly downregulated transcripts and blue dots represent significantly upregulated transcripts in *SETDB1*, and *wde* GKD ovaries compared with control ovaries (FDR = False Discovery Rate < 0.05 and genes with 1.5-fold or higher change were considered significant). (C) Venn diagram of upregulated genes from RNA-seq of *SETDB1* and *wde* GKD ovaries compared to *UAS-Dcr2;NG4NGT*. 862 targets are shared between GKD of *SETDB1* and *wde*, suggesting that SETDB1 and Wde co-regulate a specific cohort of genes. (D) The most significant biological process GO terms of shared upregulated genes in ovaries depleted of *SETDB1* and *wde* compared to *UAS-Dcr2;NG4NGT* control (FDR from p-values using a Fisher’s exact test), showing differentiation as one of the significant processes regulated by SETDB1/Wde. (E-F) RNA-seq track showing that *RpS19b* and *blanks* is upregulated upon GKD of *SETDB1* and *wde*. (G) Violin plot of mRNA levels of the 862 shared upregulated targets in ovaries enriched for GSCs, cystoblasts, cysts, and whole ovaries, showing that the shared targets between *SETDB1* and *wde* are most highly expressed up to the cyst stages, that then tapers off in whole ovaries. Statistical analysis performed with Hypergeometric test; *** indicates p<0.001. (H-J1) Confocal images of germaria probed for *RpS19b* mRNA (red, grayscale) and DAPI (blue) in *UAS-Dcr2;NG4NGT* (H-H1) showing *RpS19b* RNA expression restricted to early stages of oogenesis and in GKD of *SETDB1* (I-I1) and *wde* (J-J1) ovarioles showing *RpS19b* mRNA expression is expanded to egg chambers. (K-M1) Confocal images of germaria probed for *blanks* mRNA (red, grayscale) and DAPI (blue) in *UAS-Dcr2;NG4NGT* (K-K1) showing *blanks* mRNA expression restricted to early stages of oogenesis and in GKD of *SETDB1* (L-L1) and *wde* (M-M1) ovarioles where *blanks* mRNA expression is expanded to egg chambers. (N-O) Quantification of fluorescence intensity of *RpS19b* (N) and *blanks* (O) mRNAs in the germarium and egg chambers during development in ovaries depleted of *SETDB1* (magenta) or *wde* (orange) compared to control ovaries (gray). Statistical analysis was performed with Dunnett’s multiple comparisons test; N= 10 ovarioles; ns = p>0.05, *=p<0.05, ** = p<0.01, *** = p<0.001.

SETDB1 and Wde are known to repress gene expression, thus we first focused on mRNAs with increased levels in the GKD ovaries (Clough et al., 2014; Osumi et al., 2019). Gene Ontology (GO) analysis of the shared upregulated RNAs indicated that many were genes involved in differentiation (**Figure 2D)**. Among the upregulated RNAs was *RpS19b*, validating our initial screen, as well as genes that promote synaptonemal complex formation such as *sunn, ord* and *cona* (**Figure 2E; Figure S2C-E**). In addition, the *blanks* mRNA, which is highly expressed only in GSCs, cystoblasts and early cysts of WT, was upregulated and ectopically expressed in the egg chambers of *SETDB1-* and *wde-*GKD ovaries (**Figure 2F; Figure S2F-I**) (Blatt et al., 2021). Blanks is a component of a nuclear siRNA pathway that has critical roles in the testis but does not have any overt function during oogenesis (Gerbasi et al., 2011). Thus, *SETDB1* and *wde* repress a cohort of RNAs that are either critical for transition from GSC to an oocyte or merely expressed during early oogenesis.

To determine when during oogenesis SETDB1 and Wde act to repress genes, we analyzed available RNA-seq libraries that were enriched for GSCs, cystoblasts, and cysts, early egg chambers and late-stage egg chambers (McCarthy et al., 2019). We found that *SETDB1*/*wde*-regulated RNAs decreased after the cyst stages and their levels were attenuated in the later stages of oogenesis compared to non-targets (**Figure 2G, Figure S2J-L)** (McCarthy et al., 2019). This reduction did not happen in absence of *SETDB1* and *wde* (**Figure 2G**). RNA *in situ* analysis of *blanks*, and *RpS19b* revealed that these mRNAs are present in the early stages of oogenesis and are attenuated after oocyte specification in controls but that these RNAs persisted in *SETDB1* and *wde* GKD egg chambers (**Figure 2H-O**). Thus, mRNAs that are broadly expressed prior to oocyte specification, become repressed by SETDB1 and Wde in differentiated egg chambers.

### SETDB1 represses transcription of a subset of targets by increasing H3K9me3 enrichment

To investigate whether SETDB1/Wde-regulated mRNAs are repressed at the level of transcription, we examined a subset of nascent mRNAs (pre-mRNAs) by qRT-PCR. Indeed, the levels of nascent *RpS19b*, *ord*, *sunn*, *cona* and *blanks* mRNAs were increased in *SETDB1*/*wde-* GKDs ovaries compared to control WT ovaries (**Figure S3A-B**). These data suggest that transcription of these genes increases upon loss of SETDB1 or Wde.

To determine if the SETDB1-dependent repression of these genes involves changes in H3K9me3, we performed CUT&RUN (Ahmad, 2018; Skene and Henikoff, 2017) on adult WT ovaries enriched for differentiated egg chambers where these genes are repressed (**Figure 2G**). Analysis of CUT& RUN data from adult WT showed enrichment of H3K9me3 marks on previously identified SETDB1 targets and genes containing heterochromatin such as PHD Finger Protein 7 (*phf7)* and *light* (*lt)* respectively validating our CUT&RUN data (**Figure 3A-B; Figure S3C)** (Devlin et al., 1990; Smolko et al., 2018). As genes in *Drosophila* genome are closely packed, we only analyzed the gene body from 5’UTR to the end of the 3’UTR to unambiguously identify SETDB1 regulated genes (Schwartz and Cavalli, 2017). We found that 1593 out of 2,316 genes upregulated upon loss of SETDB1 are enriched for H3K9me3 marks compared to IgG negative control (**Figure 3C**). In addition, we found that 888 genes lose H3K9me3 on their gene bodies upon GKD of *SETDB1* including *RpS19b* and ATP-dependent chromatin assembly factor (*Acf)* (**Figure 3D-F**). The upregulated genes that do not show changes to H3K9me3 marks within the gene body may be regulated by elements outside of the gene body or indirectly. Importantly, taken together, our data suggest that SETDB1 is required for H3K9me3 enrichment and transcriptional repression of a cohort of early-oogenesis genes in the egg chamber.

**Figure 3:**
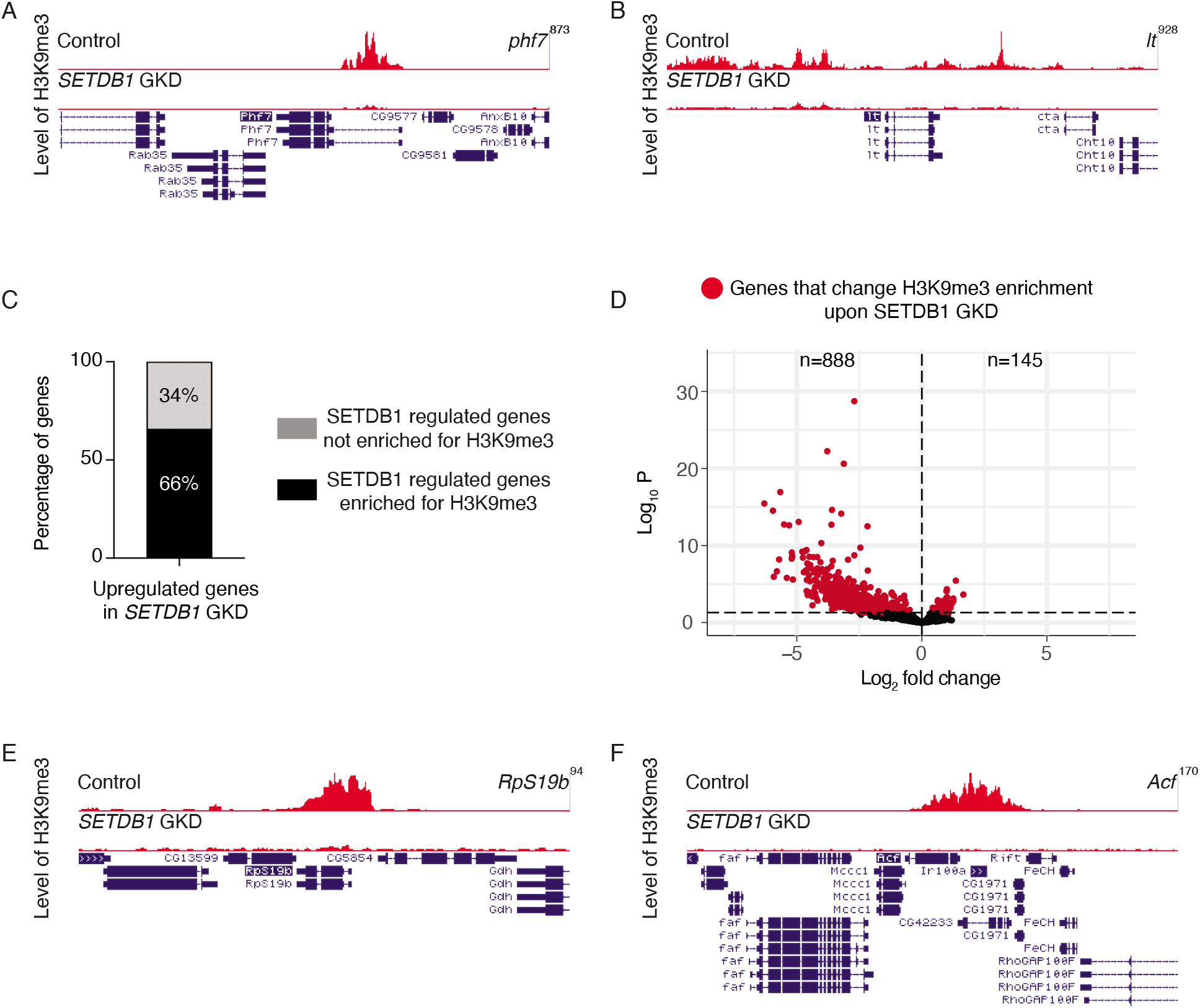
SETDB1 promotes silencing of early oogenesis genes by regulating levels of H3K9me3. (A-B) Tracks showing level of H3K9me3 on previously validated and known heterochromatic genes *phf7* and *lt* respectively. (C) Bar graph showing genes regulated by *SETDB1* that are enriched for H3K9me3 on the gene body. 1593 (black) out of 2316 (gray) genes upregulated upon loss of *SETDB1* are enriched for H3K9me3. (D) Volcano plot showing changes in H3K9me3 in *SETDB1* GKD compared to WT. Genes that lose H3K9me3 are shown on the left (red). 888 genes lose H3K9me3 after *SETDB1* GKD. (E-F) Tracks showing level of H3K9me3 on target genes. Our data shows loss of H3K9me3 on SETDB1 targets *RpS19b* and *Acf* respectively (E-F) suggesting they are directly regulated by SETDB1.

*SETDB1* is required for transposon repression during oogenesis (Andersen et al., 2017; Rangan et al., 2011), and the upregulation of transposons can affect gene expression (Sienski et al., 2012; Upadhyay et al., 2016). However, we found that the upregulation of genes in the differentiated stages that we observed upon depletion of *SETDB1* was not due to the secondary effect of transposon upregulation as the expression of *RpS19b* reporter was not altered in germline depleted of *aubergine* (*aub*), a critical component of the piRNA pathway (**Figure S3D-F**) (Chen et al., 2007; Czech et al., 2018; Malone et al., 2009; Wang et al., 2015). Nor, did *aub* depletion cause mid-oogenesis death as we observed in *SETDB1* and *wde* GKDs (**Figure S3D-F**) (Chen et al., 2007; Wilson et al., 1996). Overall, our data suggest that loss of *SETDB1* derepresses a subset of genes during late oogenesis via decreased H3K9me3, independent of transposon dysregulation.

### SETDB1 is required for the expression of NPC components

GO term analysis of downregulated targets of *SETDB1*/*wde* GKD included genes that regulate transposition, consistent with the previously described role of SETDB1/Wde in the piRNA pathway and those that regulate proper oocyte development, consistent with the previously described phenotype (**Figure 4A**)(Andersen et al., 2017; Clough et al., 2007; Koch et al., 2009; Rangan et al., 2011).

**Figure 4:**
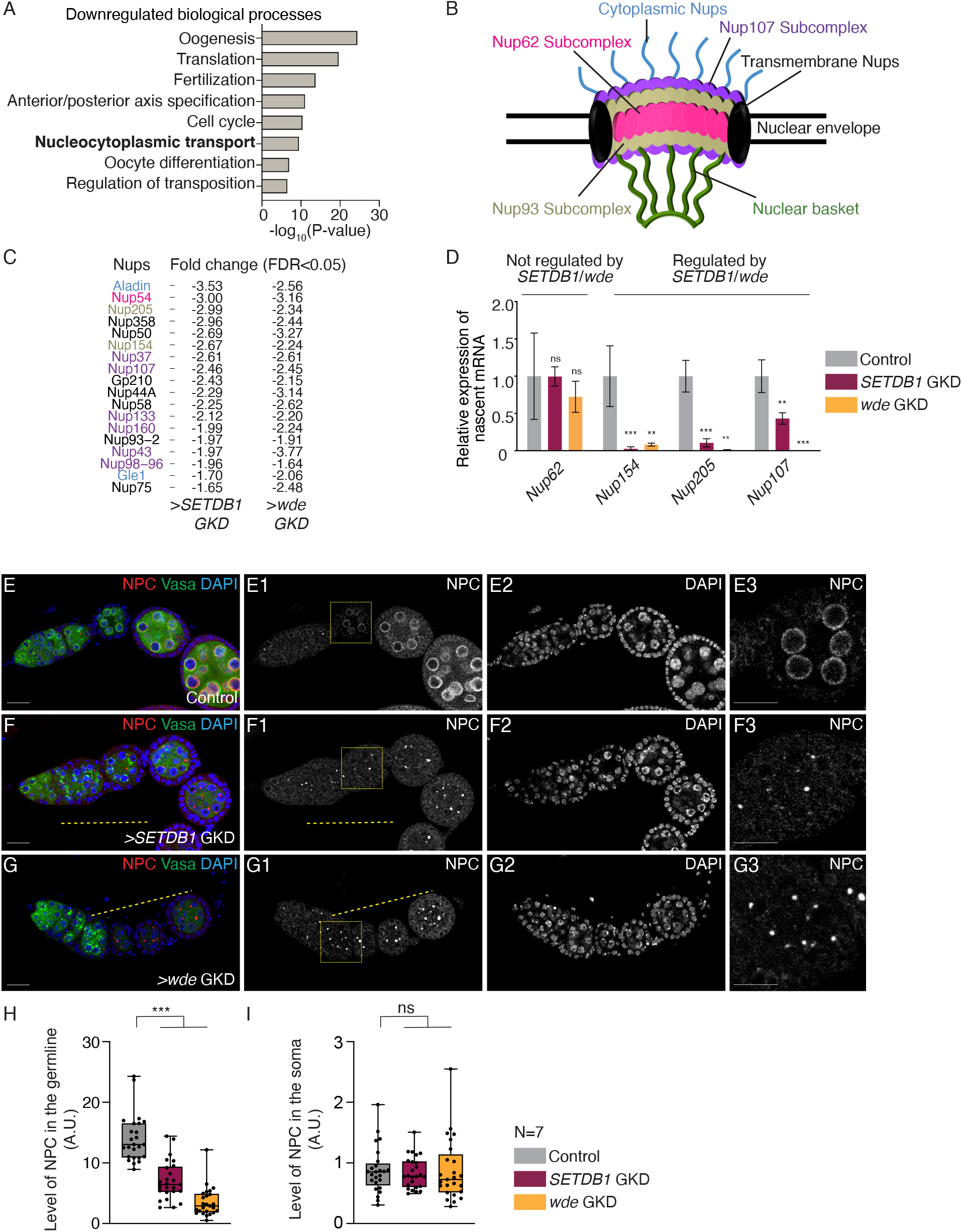
SETDB1/Wde promotes expression of a subset of nucleoporin genes and NPC formation. (A) The significant biological process GO terms of common downregulated genes in *SETDB1* or *wde* GKD ovaries compared to *UAS-Dcr2;NG4NGT* control (FDR from p-values using a Fisher’s exact test), showing nucleocytoplasmic transport as one of the significant processes regulated by SETDB1/Wde. (B) A schematic of the Nucleopore Complex (NPC) showing cytoplasmic ring, nuclear ring and basket facing nucleoplasm and a central scaffold spanning the nuclear membrane. NPC is composed of several subcomplexes and ~30 nucleoporins (nups). (C) Table showing levels of 18 nucleoporin mRNAs that are down regulated 1.5 or more fold in both *SETDB1* or *wde* GKD ovaries compared to *UAS-Dcr2;NG4NGT* control ovaries. (D) qRT-PCR assaying the pre-mRNA levels of *SETDB1* and *Wde*-regulated *Nup* genes, including *Nup154*, *Nup205* and *Nup107* are decreased compared to control *UAS-Dcr2;NG4NGT* while levels of non-target *Nup62* pre-mRNA is not affected (control level vs *SETDB1* GKD and *wde* RNA in=3, ** = p<0.01, *** = p<0.001, Error bars are SEM, Student’s t-Test). (E-G3) Ovariole and egg chamber images of control *UAS-Dcr2;NG4NGT* (E-E3), GKD of *SETDB1* (F-F3) and *wde* (G-G3) stained for NPC (red, grayscale), Vasa (green) and DAPI (blue). NPC staining was done using mab414 antibody. Depletion of *SETDB1* shows reduced expression of NPC in the egg chambers suggesting SETDB1 regulates expression of several nucleoporins which in turn regulates formation of NPC. (H-I) A.U. quantification of NPC level in the germline (H) and soma (I) in *SETDB1* and *wde* GKD ovaries compared to *UAS-Dcr2;NG4NGT* control. Statistical analysis was performed with Dunnett’s multiple comparisons test; N= 25 ovariole for germline and 15 for somatic quantitation; ns = p>0.05, * = p ≤ 0.05, ** = p<0.01, *** = p<0.001.

Unexpectedly, we observed that genes involved in nucleocytoplasmic transport were downregulated in *SETDB1/wde*-GKD ovaries as compared to controls (**Figure 4A**). Nucleocytoplasmic transport is mediated by Nucleopore complexes (NPCs), which span the nuclear membrane and consist of a cytoplasmic ring, a central scaffold spanning the nuclear envelope, and a nuclear ring and basket (**Figure 4B**) (M. Capelson et al., 2010; Doucet and Hetzer, 2010; Gozalo and Capelson, 2016). Beyond regulating nucleocytoplasmic transport, NPCs also regulate gene transcription, for instance by anchoring and maintaining heterochromatic domains (Capelson and Hetzer, 2009; Hou and Corces, 2010; Iglesias et al., 2020; Sarma and Willis, 2012; Sood and Brickner, 2014). We found that GKD of *SETDB1*/*wde* in the germline resulted in downregulation of 18 out of ~30 nucleoporins (Nups) that make up the Nucleopore complex in *Drosophila* (**Figure 4C**), including a germline enriched *Nup154* that is critical for oogenesis (Colozza et al., 2011; Gigliotti et al., 1998; Grimaldi et al., 2007). The Nups that were downregulated upon depletion of *SETDB1* and *wde* were not isolated to one specific NPC subcomplex (**Figure 4B-C**).

We found that nascent mRNAs corresponding to the *SETDB1/Wde* targets *Nup154*, *Nup205* and *Nup107* were downregulated in *SETDB1*/*wde*-GKD ovaries, whereas the non-target *Nup62* was unaffected, suggesting that SETDB1/Wde promotes transcription of a cohort of *Nups* (**Figure 4D**). In addition, the levels of a Nup107-RFP fusion protein, under endogenous control (Katsani et al., 2008), were significantly reduced in the cysts and egg chambers of *SETDB1-* and *wde-*GKD compared to controls (**Figure S4A-D**).

To determine if loss of *Nup* expression in *SETDB1*/*wde-*GKD ovaries resulted in loss of NPC formation, we performed immunofluorescence with an antibody that is known to mark NPCs in *Drosophila* (Maya Capelson et al., 2010; Davis and Blobel, 1987; Hampoelz et al., 2019; Kuhn et al., 2019). We found that NPC levels were reduced in the egg chambers of *SETDB1*/*wde-*GKD ovaries compared to controls (**Figure 4E-H**), but the nuclear lamina was unaffected (**Figure S4E-H**), and NPCs in the soma were also unaffected (**Figure 4I**). Thus, *SETDB1*/*wde* are required for the proper expression of Nups and NPC formation after oocyte specification.

Heterochromatic genes and piRNA clusters require heterochromatin to promote their transcription (Rangan et al., 2011; Weiler and Wakimoto, 1995). Although we found that *SETDB1* is required for upregulation of Nups, CUT&RUN analysis of H3K9me3 marks revealed that only 3 of the *Nup* genes had any enrichment of H3K9me3 (*Mbo*, *Nup188*, *Gp210*). Moreover, among *SETDB1*-regulated *Nups*, only Gp210 showed any heterochromatic enrichment **(Supplemental Table 2).** Taken together, we find that *SETDB1* promotes proper expression of *Nups* by a yet unknown mechanism in the germline.

### Nucleoporins are required to maintain heterochromatin domains at the nuclear periphery

Our data so far indicate that, in *Drosophila* female germline, heterochromatin formation mediated by SETDB1 is required for proper NPC formation by promoting proper expression of a subset of *Nups* including *Nup107* and *Nup154* (**Figure 4C**). In yeast, a subset of Nups are part of the heterochromatin proteome and are required to cluster and maintain heterochromatin at the NPC (Iglesias et al., 2020). This subset includes Nup107 and the yeast homolog of Nup154, Nup155, which both have reduced expression in *SETDB1/Wde*-GKD compared to controls. We hypothesized that in *Drosophila*, SETDB1 could promote silencing of early oogenesis genes by promoting heterochromatin formation. This heterochromatin then promotes expression of Nups and NPC formation, which can then help maintain heterochromatin by anchoring it to nuclear periphery and thus promoting silencing of early-oogenesis genes.

To first determine if heterochromatin and nucleoporins associate in *Drosophila* female germline, we utilized antibody against H3K9me3 to mark heterochromatin and Nup107-RFP to mark NPCs in WT ovarioles (Katsani et al., 2008; Rangan et al., 2011). We found that H3K9me3 domains were often at the nuclear periphery, in close proximity with Nup107-RFP (**Figure 5A-A2, E**). Next, to determine if loss of *Nups* leads to loss of heterochromatin, we first depleted *Nup154* and probed for heterochromatin formation. We chose *Nup154*, as its loss of function phenotype of *Nup154* has been well described (Gigliotti et al., 1998; Grimaldi et al., 2007). We found that GKD of *Nup154* in the germline, resulted in egg chambers that do not grow and die mid-oogenesis as previously described for *Nup154* mutants **(Figure S5A-B2)** (Gigliotti et al., 1998). In addition, depletion of *Nup154* results in proper translocation of SETDB1 from the cytoplasm to the nucleus suggesting that transport of SETDB1 into the nucleus is not grossly affected **(Figure S5C-D1)**. By staining for H3K9me3 marks, we found that upon GKD of *Nup154*, heterochromatin domains initially form (**Figure S5E-F2)**. However, in the egg chambers of *Nup 154* GKD, the colocalization between H3K9me3 domains and Nup107-RFP levels at the nuclear periphery were significantly reduced prior to significant reduction of heterochromatin levels **(Figure 5A-D1**, **Figure S5E-F3, I)**. GKD of *Nup107* also resulted in egg chambers that do not grow and loss of heterochromatin **(Figure S5E-I)**. Thus, *Nups 154* and *107*, which are positively regulated by *SETDB1*, are required for H3K9me3 localization at the nuclear periphery for H3K9me3 maintenance in the female germline.

**Figure 5:**
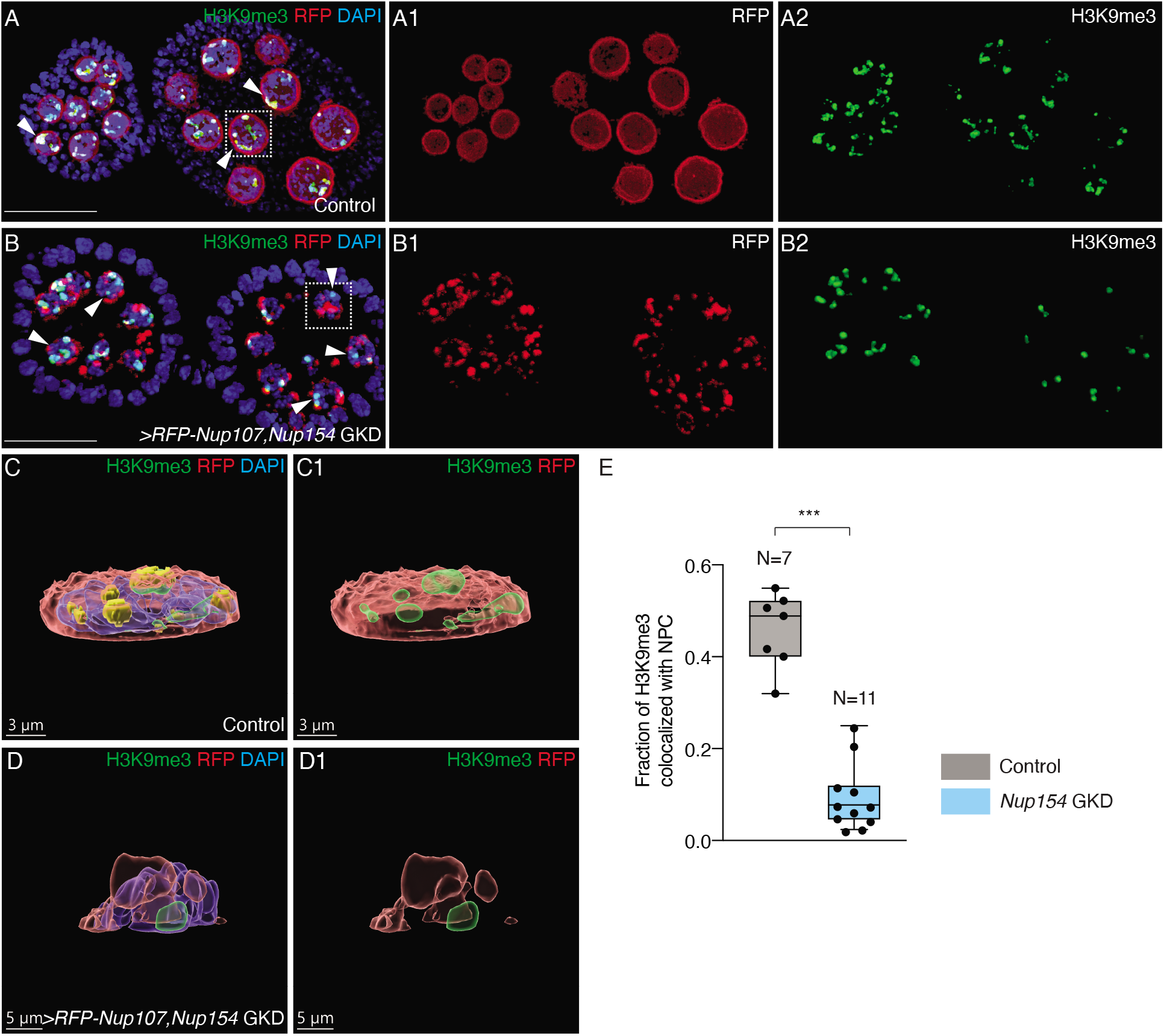
H3K9me3 heterochromatin colocalizes with NPC component Nup107 at the nuclear periphery. (A-A2) Egg chambers of control *UAS-Dcr2;NG4NGT* ovariole showing RFP-Nup107 (red, right red channel), H3K9me3 (green, right green channel). Heterochromatin is seen in close association with NPC (white arrows). Colocalized fraction is shown in yellow. (B-B2) Egg chambers of *Nup154* GKD ovariole showing significant decrease in the colocalization (white arrows) between RFP-Nup107 (red, right red channel) and H3K9me3 (green, right green channel). (C-C1) 3D reconstruction of a single nuclei (white dotted box in A) from an egg chamber of control *UAS-Dcr2;NG4NGT* ovariole. Yellow channel shows colocalized fraction of RFP-Nup107 (red) and H3K9me3 (green). This shows that H3K9me3 heterochromatin domains (green) are formed at the nuclear periphery and closely associate with Nup107. (D-D1) 3D reconstruction of a single nuclei (white dotted box in B) from an egg chamber of *Nup154* GKD showing colocalization (yellow) of H3K9me3 with Nup107. This shows significant reduction in colocalized fraction of H3K9me3 with Nup107. (E) Quantification of levels of H3K9me3 that colocalizes with NPC in the germline of control ovarioles (gray) in contrast to *Nup154* GKD ovarioles (blue). Quantitative object based colocalization was measured in Imaris software, *** = p<0.001, one-tailed Students t-Test.

### Nups are required for silencing early-oogenesis genes

Based on our findings above that Nups are required to maintain H3K9me3 levels and localization, we hypothesized that they are also required to silence the early-oogenesis RNAs in differentiated egg chambers. To test this hypothesis, we depleted *Nup154* and *Nup107* in the germline of a fly carrying the *RpS19b-GFP* reporter. We found that GKD of these nucleoporins resulted in upregulation of RpS19b-GFP phenocopying GKD of *SETDB1/wde* (**Figure 6A-C; Figure S6A-B1, D**). Moreover, germline depletion of *Nup62*, which is within the NPC but not regulated by SETDB1, also resulted in upregulation of *RpS19bGFP* and egg chambers that did not grow (**Figure S6A-D**). This suggests that activity of NPC components and not just the Nups regulated by SETDB1 are required for silencing *RpS19b-GFP* reporter.

**Figure 6:**
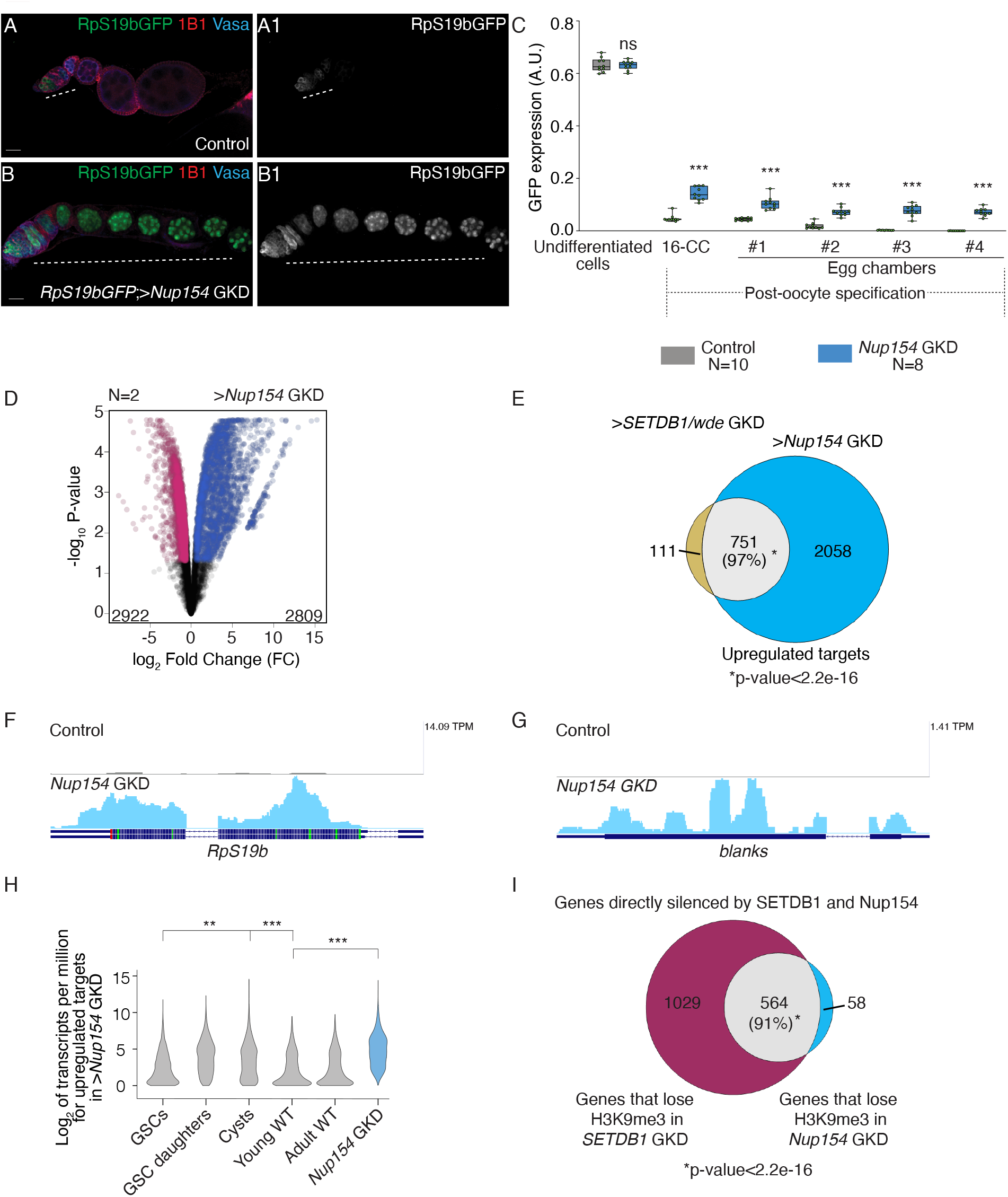
Nup154 is required for silencing a cohort of genes expressed during early oogenesis. (A-B1) Ovariole of control *RpS19b-GFP* (A-A1), GKD of *Nup154* (B-B1) stained for GFP (green, right grayscale), Vasa (blue) and 1B1 (red). Depletion of *Nup154* shows characteristic phenotype where the egg chambers did not grow and there was ectopic expression of *RpS19b-*GFP in the egg chambers (white dashed line). (C) Arbitrary units (A.U.) quantification of RpS19b-GFP expression in the germarium and egg chambers during development upon GKD of *Nup154* (blue) compared to control ovaries (gray). GFP is expressed higher in single cells in the germarium, decreases in the cyst stages, and then attenuated upon egg chamber formation in control. In *Nup154* GKD, GFP expression persists in the egg chambers. Statistical analysis was performed with Dunnett’s multiple comparisons test; N= 10 and 8 ovarioles for control and *Nup154* GKD respectively; ns = p>0.05, *=p<0.05, ** = p<0.01, *** = p<0.001. (D) Volcano plots of –Log_10_P-value vs. Log2Fold Change (FC) of mRNAs that show changes in *Nup154* GKD compared to *UAS-Dcr2;NG4NGT* control ovaries. Pink dots represent significantly downregulated transcripts and blue dots represent significantly upregulated transcripts in *Nup154* GKD ovaries compared with control ovaries (FDR = False Discovery Rate < 0.05 and 1.5-fold or higher change were considered significant). (E) Venn diagram of upregulated overlapping genes from RNA-seq of *SETDB1* and *wde* and genes from *Nup154* germline depleted ovaries compared to *UAS-Dcr2;NG4NGT*. 751 upregulated targets are shared between *SETDB1, wde* and *Nup154* GKD, suggesting that *Nup154* and *SETDB1* function in co-regulating a specific set of genes. (F-G) RNA-seq track showing that *RpS19b* (F) and *blanks* (G) are upregulated upon germline depletion of *Nup154*. (H) Violin plot of mRNA levels of the 2809 upregulated targets in ovaries enriched for GSCs, cystoblasts, cysts, and whole ovaries, showing that the upregulated targets of *Nup154* are most highly enriched upto the cyst stages, and then tapers off in whole ovaries. Statistical analysis performed with Hypergeometric test; *** indicates p<0.001. (I) Venn diagram showing overlapping genes that lose H3K9me3 after depletion of both *SETDB1* and *Nup154* in the germline. 622 genes lose H3K9me3 after *Nup154* GKD out of which 564 genes are also directly silenced by SETDB1, suggesting co-regulation of these genes by both SETDB1 and Nup154.

To determine if Nups are required for silencing other early oogenesis RNAs, we performed RNA-seq, and compared *Nup154* GKD ovaries with young ovaries as a developmental control **(Figure S2A)**. Using a 1.5-fold cut off (Fold Change (FC)≥|1.5|) and False discovery rate (FDR)<0.05), we found that compared to control, in *Nup154* GKD 2809 genes are upregulated, and 2922 genes are downregulated (**Figure 6D) (Supplemental Table 1)**. Strikingly, 97% of upregulated genes and 89% of downregulated *SETDB1*/*Wde* targets overlapped with *Nup154* GKD (**Figure 6E; S6E**). *Nup154* was involved in silencing genes that promote oocyte differentiation including synaptonemal complex components *ord*, *sunn* and *cona* as well as *RpS19b* (**Figure 6F; S6F-H**). In addition, GKD of *Nup154* also resulted in upregulation of *blanks* (**Figure 6G**). The levels of *Nup154*-regulated RNAs decreased after the cyst stage, when the oocyte is specified, in contrast to non-targets, which have similar RNA levels at all stages (**Figure 6H; S6I-J**). Thus, Nup154 is critical for silencing early-oogenic mRNAs in the differentiated egg chambers.

To determine if *Nup154* is required for H3K9me3 marks at SETDB1-regulated gene, loci such as *RpS19b*, we carried out CUT& RUN for H3K9me3 in control and *Nup154* GKD. We found that 564 out of 622 genes displaying a loss in H3K9me3 in Nup154 GKD also show the same loss in *SETDB1* GKD including *RpS19b* and *Acf* (**Figure 6I; S6K-L**) **(Supplemental Table 2)**. Taken together, we find that Nups are required for silencing and H3K9me3 at a subset of *SETDB1/Wde-*regulated loci.

### Silencing genes expressed during the early oogenesis stages is required for maintaining oocyte fate

We next asked why loss of *SETDB1, wde* and *Nups* results in egg chambers that do not grow and die mid-oogenesis. Egg chambers with oocyte specification or maintenance defects result in death of egg chambers mid-oogenesis (Blatt et al., 2021). To determine if there are oocyte specification or maintenance defects, we stained GKD of *SETDB1, wde* and *Nup154* for the oocyte marker Egalitarian (Egl) as well as Vasa and 1B1 (Mach and Lehmann, 1997; Navarro et al., 2004). In the early stages of oogenesis, as in control, GKD of *SETDB1, wde* and *Nup154* resulted in one Egl positive cell, suggesting that oocyte is specified (**Figure 7A-E**). While initial Egl localization to oocytes appeared to be normal, we cannot rule out subtle specification defects. However, in the later egg chambers, compared to control ovariole, GKD of *SETDB1, wde* and *Nup154* resulted in either mis-localization or diffuse Egl expression suggesting loss of oocyte fate (**Figure 7A-E**). Taken together, these data suggest that SETDB1, Wde and Nup154 are required for maintaining the oocyte fate.

**Figure 7:**
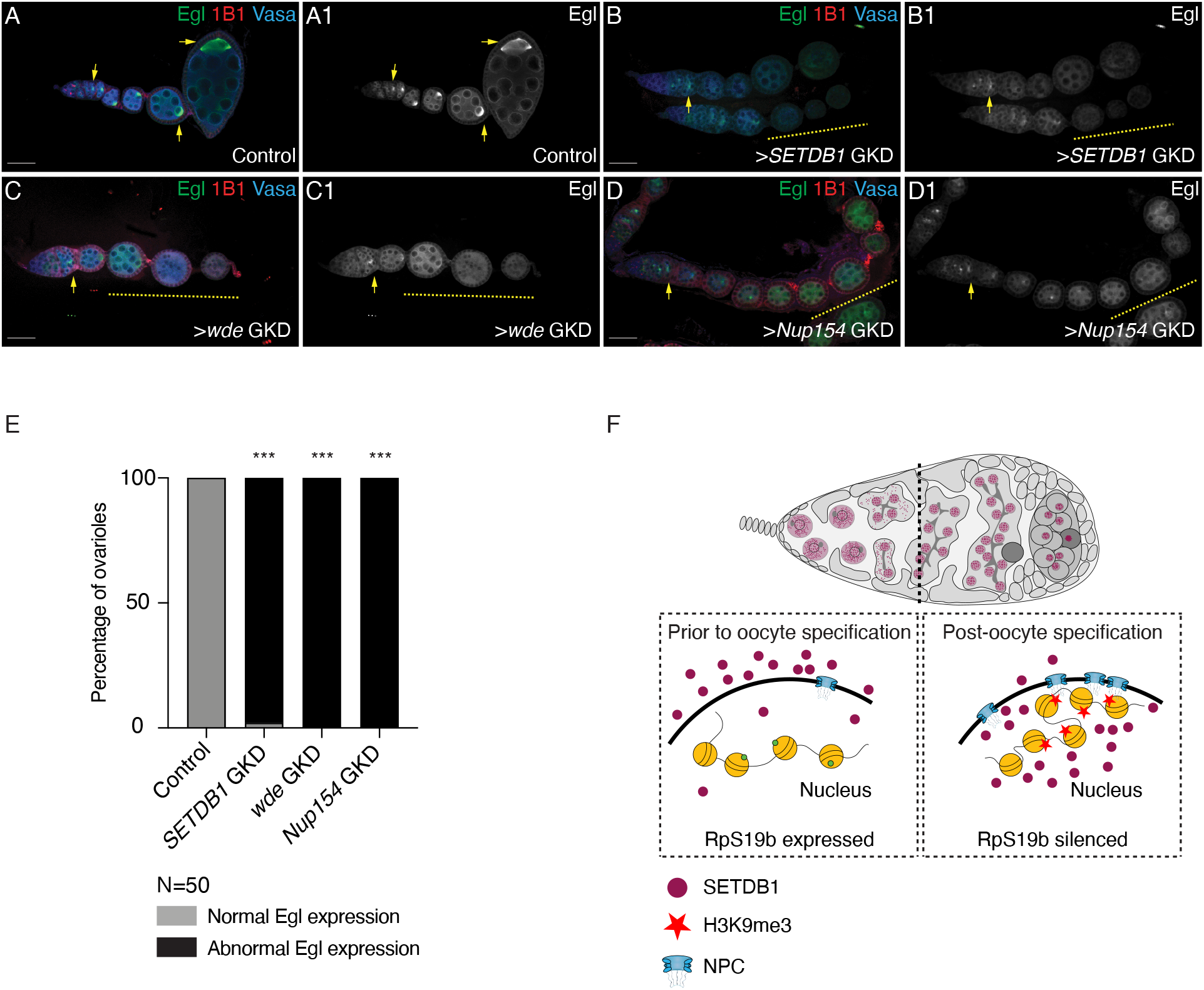
Silencing of early oogenesis genes mediated by SETDB1, Wde and Nup154 is required for maintenance of oocyte fate. (A-D1) Ovarioles of control *UAS-Dcr2;NG4NGT* (A-A1), GKD of *SETDB1* (B-B1), *wde* (C-C1) and *Nup154* (D-D2) stained for Egl (green, right grayscale), Vasa (blue) and 1B1 (red). Control shows proper oocyte specification with one oocyte in each egg chamber. Depletion of *SETDB1*, *wde* and *Nup154* in the germline results in initial oocyte specification (yellow arrow) which is then lost in the subsequent egg chambers (yellow dashed line). (E) Quantification of percentage ovarioles with abnormal/loss of Egl expression (black) in ovaries depleted of *SETDB1* or *wde* or *Nup154* compared to control ovaries (gray) (N= 50 ovarioles; 98% in *SETDB1* GKD and 100% in *wde* and *Nup154* GKD compared to 0% in control.) Statistical analysis was performed with Fisher’s exact *** = p<0.001. (F) A model showing that nuclear translocation of SETDB1 after differentiation promotes heterochromatin formation mediated by deposition of H3K9me3 mark. This heterochromatin promotes increased NPC formation which then helps maintain heterochromatin.

## Discussion

Many maternally contributed mRNAs in oocytes are critical for early development after fertilization (Calvi et al., 1998; Huynh and St Johnston, 2004; Kugler and Lasko, 2009; Navarro et al., 2004; Telfer, 1975). We previously showed that many mRNAs expressed in germ cells and the undifferentiated stages of oogenesis must be selectively degraded and thus excluded from the maternal contribution (Blatt et al., 2021). However, the potential role of transcriptional silencing of germ cell and GSC-enriched genes during oogenesis was unclear. Here, we found that regulated translocation of SETDB1 into the nucleus during oocyte specification is required to silence germ cell- and early oogenesis-genes in the differentiated egg chambers (**Figure 7F**), and that this process is essential to maintain oocyte fate. Thus, some genes that are expressed in germ cells and some that promote differentiation are transcriptionally silenced at the onset of oocyte specification mediated by a feedback loop between heterochromatin and NPC.

### Regulated heterochromatin formation during oocyte specification promotes germ cell to oocyte transition

A large fraction of SETDB1 is cytoplasmic in the undifferentiated stages of the germline. As the oocyte is being specified during differentiation, SETDB1 becomes mostly nuclear (Clough et al., 2014). This translocation of SETDB1 to the nucleus during oocyte specification is mediated by Windei (Wde), the *Drosophila* ortholog of mAM/MCAF1 (Koch et al., 2009; Osumi et al., 2019). Here we find that translocation of SETDB1 to the nucleus during oocyte specification is required to silence germ cell and early-oogenesis genes at the onset of oocyte specification. MCAF1 also regulates the accumulation of SETDB1 in the nucleus in mammalian cells (Tsusaka et al., 2019). In addition, loss of *SETDB1* during mammalian oogenesis results in meiotic defects and infertility (Eymery et al., 2016). These data suggest that regulated heterochromatin formation is conserved to promote proper oogenesis in mammals.

We discovered that SETDB1 is required to silence two major classes of genes. The first group is involved in GSC differentiation into an oocyte, including critical genes that promote meiosis I. The second group of genes are those that are merely expressed in the germ cells prior to differentiation into an oocyte, but have no specific function in the female germline such as *blanks* (Blatt et al., 2021; Gerbasi et al., 2011). We propose that these genes that are silenced upon oocyte specification are either detrimental to late oogenesis or early embryogenesis. Indeed, it has been shown that overexpression of one such gene *actin 57B* (*act57B)*, which is repressed by SETDB1/Wde **(Supplemental Table 1)**, is detrimental to oogenesis (Blatt et al., 2021; Duan et al., 2020). Remarkably, some of the mRNAs encoded by genes that are transcriptionally silenced by SETDB1 during this transition are also targeted at the post-transcriptional level for degradation by members of the no go decay pathway such as *blanks* and *Act57B* (Blatt et al., 2021). Thus, our data suggests that the regulation of gene expression during the germ cell to oocyte transition reflects a two-step process: transcriptional silencing dependent on SETDB1, and post-transcriptional degradation of mRNAs to exclude a cohort of germ cell mRNAs from the maternal contribution (Blatt et al., 2021).

SETDB1 is guided to its target transposons and piRNA clusters mediated by piRNAs (Andersen et al., 2017; Czech et al., 2018; Koch et al., 2009; Osumi et al., 2019). However, our data suggests that the piRNA pathway does not play a part in silencing germ cell and early oogenesis RNAs. We find that loss of *aub* does not result in upregulation of *RpS19b*. This result is consistent with the fact that loss of *aub* and *piwi* in the germline does not result in egg chambers that do not grow (Chen et al., 2007; Wilson et al., 1996). In somatic cells of the gonad, loss of *wde* function eliminates nuclear SETDB1 signal (Osumi et al., 2019; Timms et al., 2016). However, upon depletion of Wde, SETDB1 was still ubiquitinated, a requirement for its nuclear retention (Osumi et al., 2019). This suggests that in absence of Wde, SETDB1 can translocate to the nucleus but cannot find its targets. Osumi *et al.* (2019) suggested that Wde could mediate SETDB1 recruitment to the its targets, leading to H3K9me3 deposition. In mammals, it has been shown that transcriptional factors such as the KRAB domain-containing Zinc finger proteins recruit SETDB1 to the target genes for silencing, but such transcription factors have not been identified in the female gonad (Frietze et al., 2010; Schultz et al., 2002). Thus, SETDB1 targets germ cell and early oogenesis genes for silencing independent of the piRNA pathway but through a yet undetermined mechanism, either through Wde or through yet undetermined transcription factors.

### Nucleopore complex and heterochromatin are in a feedback loop to promote gene silencing

The NPC not only mediates selective nucleo-cytoplasmic transport of macromolecules but also regulates gene expression by anchoring chromatin domains, including heterochromatin to the nuclear periphery (Maya Capelson et al., 2010; M. Capelson et al., 2010; Holla et al., 2020; Iglesias et al., 2020; Sarma and Willis, 2012). In addition, in yeast, several Nups are also part of the heterochromatin proteome suggesting that NPCs can regulate gene expression by regulating heterochromatin (Brickner et al., 2019; Iglesias et al., 2020). Consistent with these observations, we find that in the female germline of *Drosophila*, NPC and heterochromatin are closely associated. Loss of NPCs due to depletion of individual Nups results in loss of heterochromatin and subsequent upregulation of germ cell and early oogenesis genes resulting in oogenesis defects. The 97% overlap of target genes between *SETDB1*, *wde* and *Nup154* is indicative that Nups are functioning in the same pathway as SETDB1. This suggests that in the female germline, not only do NPCs associate with heterochromatin, but that NPCs also play a role in maintaining heterochromatin and gene repression during germ cell to oocyte transition.

Silencing of early oogenesis genes at the onset of oocyte specification is timed with exit from mitotic cell cycle. *Drosophila* nucleopore complex consists of ~30 different nucleoporins some of which are solubilized during early mitotic cell division (Güttinger et al., 2009; Laurell and Kutay, 2011). Nucleoporins are recruited to the chromatin in early anaphase followed by sequential reassembly of the complex (Kiseleva et al., 2001; Kutay et al., 2021). During *Drosophila* oogenesis, in the premeiotic stage, the GSC divides to eventually produce a 16-cell cyst (Huynh and St Johnston, 2004; Koch et al., 1967; Lehmann, 2012; Spradling et al., 2011). Prophase-I of meiosis begins in cysts where the oocyte is also specified (Ables, 2015; Orr-Weaver, 1995). We find that nucleoporins promote silencing of genes that are required for initiation of meiosis I such as *Rbfox1* and synaptonemal complex components *ord*, *sunn* and *cona* once the cysts have stopped dividing and the oocyte is being specified. Taken together, our data suggests a mechanism wherein after the mitotic division of cysts have ceased and meiosis I is initiated, the reassembly of NPC simultaneously promotes silencing of the genes required for the transition from mitotic GSC division to meiotic oocyte specification.

While NPC association with heterochromatin has been described, remarkably we find that loss of heterochromatin results in attenuated expression of some but not all Nup mRNAs. Heterochromatic genes and piRNA clusters require heterochromatin to promote their transcription (Andersen et al., 2017; Devlin et al., 1990; Rangan et al., 2011). However, by analyzing CUT &RUN data and previously published ChIP data of H3K9me3 marks, we found that only one Nup regulated by SETDB1 is enriched for H3K9me3 marks. Therefore, SETDB1 indirectly promotes expression of Nups.

The number of genes that need to be silenced varies based on cell types and developmental trajectory. How levels of heterochromatin are coupled to their NPC docking sites in the cell was not known. Like heterochromatin levels, the number of NPCs also varies by cell type and during differentiation (McCloskey et al., 2018). How NPC number is regulated during development was not fully understood. Our findings in the female germline suggest an elegant tuning mechanism for heterochromatin and its NPC docking sites. Heterochromatin promotes levels of NPC which then promote heterochromatin maintenance by tethering it to the nuclear periphery. We find that this loop can be developmentally regulated by controlling levels of SETDB1 in the nucleus mediated by conserved protein Wde to promote heterochromatin formation.

## Acknowledgements

We are grateful to all members of the Rangan laboratory for discussion and comments on the manuscript. We also thank Dr. Thomas Hurd and Dr. Miler Lee for their comments on the manuscript. We would like to thank Sontheimer lab for Blanks antibody, Lehmann lab for Egl antibody, Bloomington Drosophila Stock Center, Vienna Drosophila Resource Center, Transgenic GKD Project (NIH/NIGMS R01-GM084947), The BDGP Gene Disruption Project, and FlyBase for fly stocks and reagents. Furthermore, we would like to thank CFG Facility at the University at Albany (UAlbany) for performing RNA-seq analyses. P.R. is funded by NIH/NIGMS (RO1GM11177 and RO1GM135628). M.A.S is funded by NIH NIGMS R35 138120 and A.V. is funded by R01DE030927.

## Materials and Methods

### Fly lines

The following RNAi stocks were used in this study; if more than one line is listed, then both were quantitated and the first was shown in the main figure: *SETDB1* RNAi (Perrimon lab, (Rangan et al., 2011), *Wde* RNAi (Bloomington #33339), *Nup154* RNAi (Bloomington #34710), *Nup93* RNAi (VDRC #V16189), *Nup62* RNAi (Bloomington #35695), *Nup107* RNAi (Bloomington #43189), *Nup205* RNAi (VDRC #V38608).

The following tagged lines were used in this study: *dSETDB1-HA* (Bontron Lab) (Seum et al., 2007), *RpS19b-GFP* (Buszczak Lab, (McCarthy et al., 2019), mRFP-Nup107 (Bloomington #35516).

The following tissue-specific drivers and double balancer lines were used in this study: *UAS-Dcr2;nosGAL4* (Bloomington #25751), *nosGAL4;MKRS*/TM6 (Bloomington #4442), and *If*/CyO*;nosGAL4* (Lehmann Lab).

### Reagents for fly husbandry

Flies were grown at 25-29°C and dissected between 0-3 days post-eclosion.

Fly food was made using the procedures as previously described (summer/winter mix) and narrow vials (Fisherbrand Drosophila Vials; Fischer Scientific) were filled to approximately 10-12mL (Flora et al., 2018).

### Dissection and Immunostaining

Ovaries were dissected and teased apart with mounting needles in cold PBS and kept on ice. All incubation was done with nutation. Samples were fixed for 10 minutes in 5% methanol-free formaldehyde. Ovaries were washed in 0.5 mL PBT (1X PBS, 0.5% Triton X-100, 0.3% BSA) 4 times for 5 minutes each. Primary antibodies in PBT were added and incubated at 4°C nutating overnight. Samples were next washed 3 times for 5 minutes each in 0.5 mL PBT, and once in 0.5 mL PBT with 2% donkey serum (Sigma) for 15 minutes. Secondary antibodies were added in PBT with 4% donkey serum and incubated at room temperature for 3-4 hours. Samples were washed 3 times for 10 minutes each in 0.5 mL of 1X PBST (0.2% Tween 20 in 1x PBS) and incubated in Vectashield with DAPI (Vector Laboratories) for 1 hour before mounting.

The following primary antibodies were used: mouse anti-1B1 (1:20; DSHB), Rabbit anti-Vasa (1:1,000; Rangan Lab), Chicken anti-Vasa (1:1,000; Rangan Lab) (Upadhyay et al., 2016), Rabbit anti-GFP (1:2,000; abcam, ab6556), Rabbit anti-H3K9me3 (1:500; Active Motif, AB_2532132), Mouse anti-H3K27me3 (1:500; abcam, ab6002), Rabbit anti-Egl (1:1,000; Lehmann Lab), Mouse anti-NPC (1:2000; BioLegend, AB_2565026) and Rat anti-HA (1:500; Roche, 11 867 423 001). The following secondary antibodies were used: Alexa 488 (Molecular Probes), Cy3 and Cy5 (Jackson Labs) were used at a dilution of 1:500.

### Fluorescence Imaging

The tissues were visualized, and images were acquired using a Zeiss LSM-710 confocal microscope under 20X, 40X and 63X oil objective with pinhole set to 1 airy unit. All gain, laser power, and other relevant settings were kept constant for any immunostainings being compared. Image processing was done using Fiji and gain adjustment and cropping was performed in Photoshop CC 2019.

### Colocalization analysis

Confocal images of control and Nup154-RNAi mutants labeled for RFP-Nup107, H3K9me3, and DAPI were imported into Bitplane Imaris 9.6.2 for 3D reconstruction and colocalization analysis. Colocalization between RFP-Nup107 and H3K9me3 was calculated on a per egg chamber basis using the Surface-surface coloc function of Imaris and an automatic threshold detection and the surface-to-surface coloc function. The number of colocalized voxels was then normalized to the number of H3K9me3 voxels (Valm et al., 2017).

### Egg laying assays

Assays were conducted in vials with 3 control or experimental females under testing and 1 wild type control males. Crosses were set up in triplicate for both control and experimental. All flies were 1-day old post-eclosion upon setting up the experiment. Cages were maintained at 29°C and plates were changed daily for counting. Analyses were performed for 5 consecutive days. Number of eggs laid were counted and averaged. Adult flies eclosed were counted from all the vials and averaged.

### RNA isolation

Ovaries from flies were dissected in cold 1x PBS. RNA was isolated using TRIzol (Invitrogen, 15596026) (Blatt et al., 2021; McCarthy et al., 2019).

RNA was treated with DNase (TURBO DNA-free Kit, Life Technologies, AM1907), and then run on a 1% agarose gel to check integrity of the RNA.

### RNA-seq library preparation and analysis

Libraries were prepared using the Biooscientific kit. To generate mRNA enriched libraries, total RNA was treated with poly(A)tail selection beads (Bioo Scientific Corp., NOVA-512991). Manufacturer’s instructions of the NEXTflex Rapid Directional RNA-seq Kit (Bioo Scientific Corp., NOVA-5138-08) were followed, but RNA was fragmented for 13 minutes. Library quality was assessed with a Fragment Analyzer (5200 Fragment Analyzer System, AATI, Ankeny, IA, USA) following manufacturer’s instructions. Single-end mRNA sequencing (75 base pair reads) was performed on biological duplicates from each genotype on an Illumina NextSeq500 by the Center for Functional Genomics (CFG).

After quality assessment, the sequenced reads were aligned to the *Drosophila melanogaster* genome (UCSCdm6) using HISAT2 (version 2.1.0) with the RefSeq-annotated transcripts as a guide (Kim et al., 2015). Differential gene expression was assayed by DeSeq2, using a false discovery rate (FDR) of 0.05, and genes with 2-fold or higher were considered significant. The raw and unprocessed data for RNA-seq generated during this study are available at Gene Expression Omnibus (GEO) databank under accession number: XXX. GO term enrichment on differentially expressed genes was performed using Panther (Thomas et al., 2006).

### Fluorescent *in situ* hybridization

A modified *in situ* hybridization procedure for *Drosophila* ovaries was followed. Probes were designed and generated by LGC Biosearch Technologies using Stellaris® RNA FISH Probe Designer, with specificity to target base pairs of target mRNAs. Ovaries (3 pairs per sample) were dissected in RNase free 1X PBS and fixed in 1 mL of 5% formaldehyde for 10 minutes. The samples were then permeabilized in 1mL of Permeabilization Solution (PBST+1% Triton-X) rotating in RT for 1 hour. Samples were then washed in wash buffer for 5 minutes (10% deionized formamide and 10% 20x SSC in RNase-free water). Ovaries were covered and incubated overnight with 1ul of probe in hybridization solution (10% dextran sulfate, 1 mg/ml yeast tRNA, 2 mM RNaseOUT, 0.02 mg/ml BSA, 5x SSC, 10% deionized formamide, and RNase-free water) at 30°C. Samples were then washed 2 times in 1 mL wash buffer for 30 minutes and mounted in Vectashield.

### CUT&RUN assay

Ovaries from flies were dissected in ice cold 1x PBS and ovarioles were separated by teasing after dissection with mounting needles. PBS was removed and the samples were permeabilized in 1mL of Permeabilization Solution (PBST+1% Triton-X) rotating in RT for 1 hour. Samples were then incubated overnight at 4°C in primary antibody dilutions in freshly prepared BBT+ buffer (PBST + 1% BSA + 0.5 mM Spermidine + 2 mM EDTA + 1 large Roche complete EDTA-free tablets). Primary antibody was replaced with BBT+ buffer and quickly washed twice. Samples were then incubated in ~700 ng/ml of pAG-MNase in BBT+ buffer rotating for 4 hours at 25°C. Samples were then quickly washed twice in wash+ buffer (20 mM HEPES pH7.5 + 150 mM NaCl + 0.1% BSA + 0.5 mM Spermidine + 1 large Roche complete EDTA-free tablets in water). Samples were resuspended in 150 μl Wash+C (wash+ + 100 mM CaCl_2_) and incubated for 45 minutes on nutator at 4°C. The cleavage reaction was terminated by addition of 150 μl StopR (NaCl final 200 mM + EDTA final 20 mM + 100μg/mL RNaseA) and incubating the sample at 37°C for 30 minutes. Samples were then centrifuged at 16,000 x g for 5 minutes and 300 μl of the supernatant was collected for DNA discovery. To the supernatant, 2 μL 10% SDS and 2.5 μL of 20 mg /mL Proteinase K was added and incubated at 50°C for 2 hours. Half of this was kept as a backup and half was used in bead cleanup. 20 μL AmpureXP bead slurry and 280 μL MXP buffer (20% PEG8000 + 2.5 M NaCl + 10 mM MgCl2 in water) was added to the sample and mixed thoroughly followed by 15 minutes incubation at RT. The beads were separated by magnet and supernatant was discarded. The beads were carefully washed with 80% ethanol for 30 seconds, while on the magnetic stand and air dried for 2 minutes. The beads were then resuspended in 10 μL DNase free water.

### DNA seq library preparation and analysis

The samples from CUT&RUN assay were used for library preparation using NEBNext® Ultra™ DNA Library Prep Kit for Illumina® (E7645, E7103) and adaptor ligated DNA were prepared without size selection.

### CUT&Run Data Analysis

Cut&Run libraries were sequenced as paired-end 75bp reads on the Illumina NextSeq 500 at the University at Albany Center for Functional Genomics. FASTQ files were aligned to the dm6 reference genome using HISAT2 (10.1038/s41587-019-0201-4) (-X 10 -I 1000 –no-spliced-alignment, --no-discordant). Mapping statistics and data will be available from Gene Expression Omnibus. Alignment files were sorted and indexed using samtools and were subsequently used to create bigwig files for visualization with deeptools (--binSize 10)( 10.1093/nar/gkw257). Principle component analysis between samples was performed using the multiBigwigSummary and plotPCA modules from deeptools. Only gene bodies were considered and problematic genomic regions (blacklist) were removed from the analysis (10.1038/s41598-019-45839-z). Raw read counts of H3K9me3 enrichment across gene bodies was calculated using the HOMER annotateRepeats function and differential enrichment was calculated using DESeq2 (HOMER PMID:20513432, DESeq2 citation 10.1186/s13059-014-0550-8). H3K9me3 occupied genes are those with differential enrichment of H3K9me3 compared to IgG matched control conditions using DESeq2.

### Quantitative Real Time-PCR (qRT-PCR)

1 μL of cDNA from each genotype was amplified using 5μL of SYBR green Master Mix, 0.3 μL of 10μM of each reverse and forward primers in a 10 μL reaction. The thermal cycling conditions consisted of 50°C for 2 minutes, 95°C for 10 minutes, 40 cycles at 95°C for 15 seconds, and 60°C for 60 seconds. The experiments were carried out in technical triplicate and minimum 2 biological replicates for each sample. To calculate fold change in mRNA levels, comparison was done to rp49 mRNA levels which was used as the control gene. Average of the 2^ΔCt for the biological replicates was calculated. Error bars were plotted using standard error of the ratios and P-value was determined by Students t-test.

---

**Supplementary Figure1:**
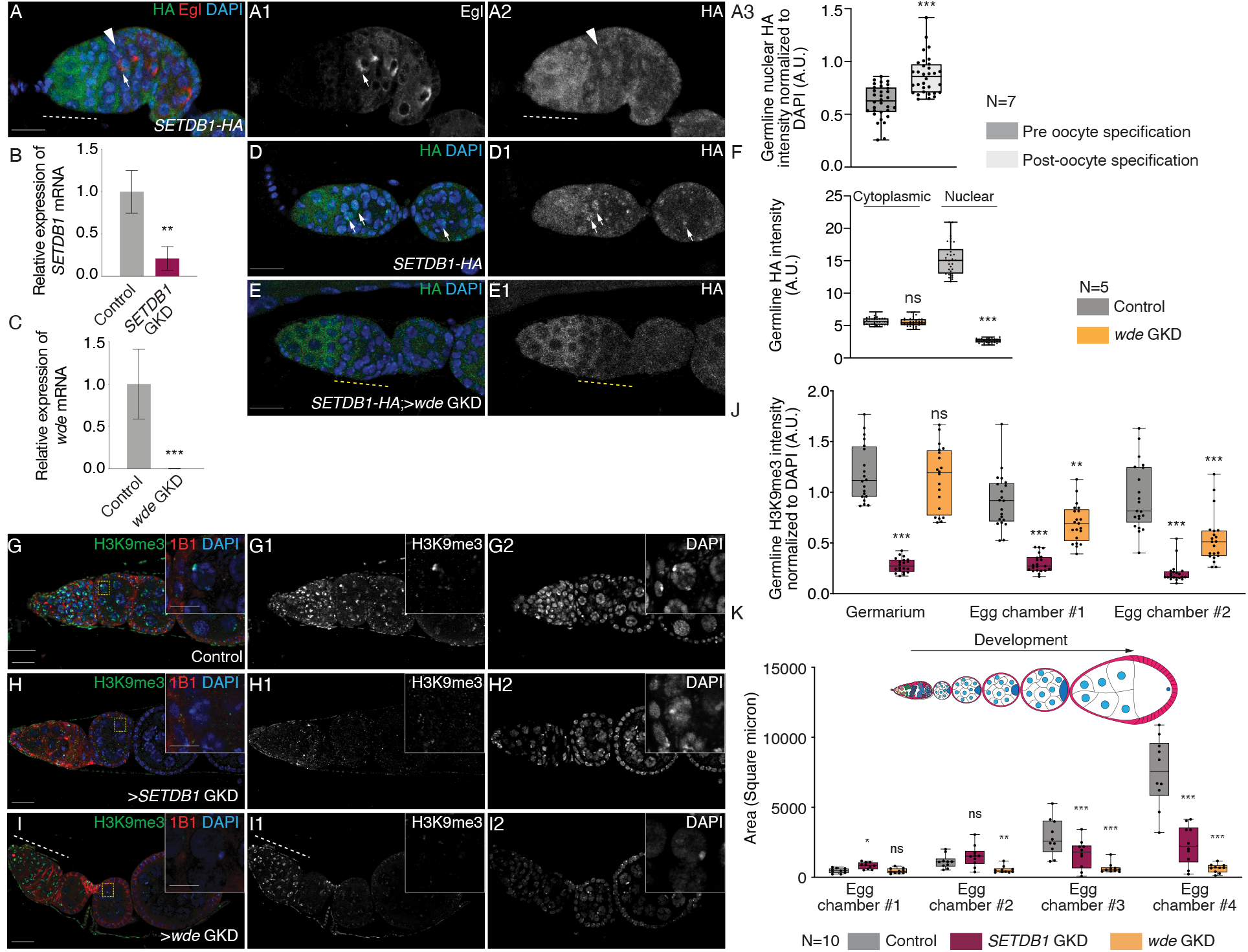
SETDB1/Wde mediated heterochromatin formation is required for silencing *RpS19b* reporter. (A-A3) Germarium of a fly carrying HA tagged SETDB1 stained for HA (green, right grayscale), oocyte marker Egl (red, right grayscale) and DAPI (blue). White arrows point at the specified oocyte. SETDB1 translocates from the cytoplasm (white dotted line) to the nucleus concurrent with oocyte specification (white solid arrow). Quantitation of HA level (A3) expressed as a ratio of nuclear SETDB1 to DAPI. Statistical analysis was performed with Welch’s t-test; N= 7 germaria; ns = p>0.05, *=p<0.05, ** = p<0.01, *** = p<0.001 (B-C) qRT-PCR assaying the mRNA levels of *SETDB1* (B) and *wde* (C) in control and *SETDB1* and *wde* GKD respectively, normalized to control *rp49* mRNA levels and indicating knockdown of these genes (N=3, **=p<0.01, *** = p<0.001, Error bars are SEM, Student’s t-test). (D-F) SETDB1-HA ovariole (D-D1) and *SETDB1-HA* ovariole depleted of *wde* (E-E1) stained for HA (green, right grayscale) and Vasa (blue). Yellow arrows point at nuclear HA. GKD of *wde* shows that levels of HA in the nucleus is attenuated (yellow dotted line). (F) Quantification of germline HA levels in the cytoplasm in the undifferentiated stages and in the nucleus of the differentiated stages in the germarium in ovaries depleted of *wde* (orange) compared to control ovaries (gray). Statistical analysis was performed with Welch’s t-test; N= 5 ovarioles; ns = p>0.05, *=p<0.05, ** = p<0.01, *** = p<0.001. (G-J) Ovariole of control *UAS-Dcr2;nosGAL4* (G-G1), GKD of *SETDB1* (H-H1) or *wde* (I-I1) stained for H3K9me3 (green, right grayscale), DAPI (blue, right grayscale) and 1B1 (red). Nurse cells from egg chamber highlighted by a dashed yellow square represent cells shown in the inset. Control shows H3K9me3 is present throughout oogenesis in the germline. GKD of *SETDB1* shows loss of H3K9me3 in all stages of the germline while depletion of *wde* results in decreased H3K9me3 post-differentiation only in the egg chambers but not in germarium (white dotted line). (J) Quantification of H3K9me3 expression in the germline normalized to DAPI level in ovaries depleted of *SETDB1* (magenta) or *wde* (orange) compared to control ovaries (gray). Statistical analysis was performed with Dunnett’s multiple comparisons test; N= 10 ovarioles; ns = p>0.05, *=p<0.05, ** = p<0.01, *** = p<0.001. (K) Quantification of area of germarium and egg chambers during development in *SETDB1* (magenta) or *wde* (orange) GKD ovaries compared to control ovaries (gray). Statistical analysis was performed with Dunnett’s multiple comparisons test; N= 10 ovarioles; ns = p>0.05, *=p<0.05, ** = p<0.01, *** = p<0.001. Scale bars are 15 micron and for main images and scale bar for insets is 4 micron.

**Supplementary Figure 2:**
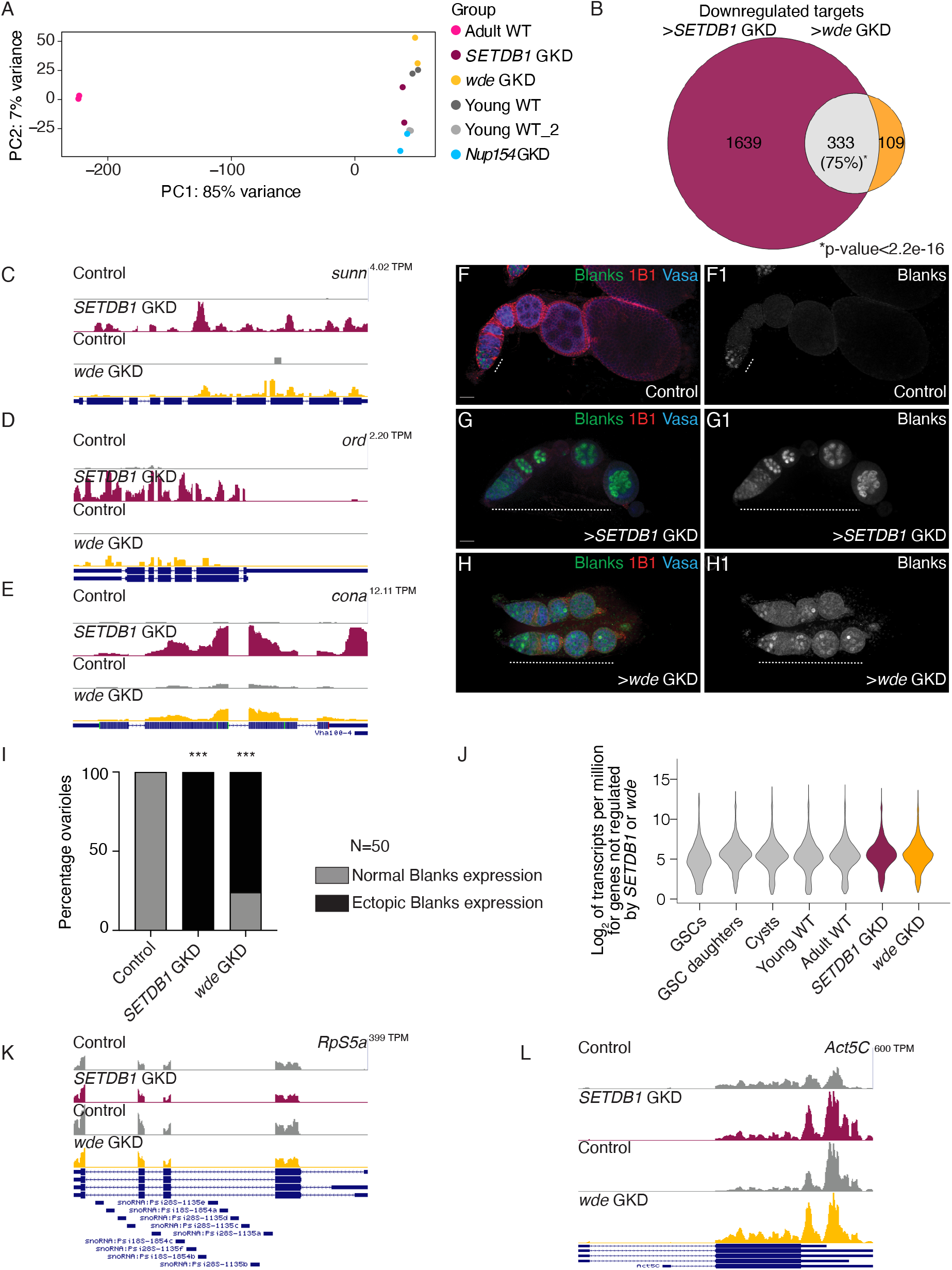
SETDB1/Wde represses a cohort of genes that are broadly expressed prior to oocyte specification. (A) Principal Component Analysis (PCA) comparing RNA-seq data sets including adult WT, young WT, *SETDB1* GKD and *wde* GKD indicates that the *SETDB1, wde* and *Nup154* GKD samples are similar to young WT. (B) Venn diagram of downregulated genes from RNA-seq of *SETDB1* and *wde* GKD ovaries compared to *UAS-Dcr2;NG4NGT*. 333 targets are shared between *SETDB1* and *wde* GKD. (C-E) RNA-seq track showing that synaptonemal complex members *sunn, ord* and *cona* are upregulated upon *SETDB1* and *wde* GKD. (F-H1) Confocal images of *UAS-Dcr2;NG4NGT* (C-C1), *SETDB1* (D-D1) and *wde* (E-E1) GKD ovarioles stained for 1B1 (red), Vasa (blue) and Blanks protein (green and grayscale) showing expanded Blanks expression in both *SETDB1* and *wde* GKD egg chambers (arrow). (I) Quantification of percentage ovarioles with ectopic *Blanks* expression (black) in *SETDB1* or *wde* GKD ovaries to control ovaries (N= 50 ovarioles; 100% in *SETDB1* GKD and 82% in *wde* GKD compared to 0% in control.) Statistical analysis was performed with Fisher’s exact test on ectopic *Blanks* expression; *** = p<0.001. (J) Violin plot of mRNA levels of the genes not regulated by *SETDB1* or *Wde* in ovaries enriched for GSCs, cystoblasts, cysts, and whole ovaries, showing that *SETDB1* and *wde* non-targets are not attenuated in the later egg chamber ovaries compared to earlier stages of oogenesis. (K-L) RNA-seq track showing that levels of non-targets *RpS5a* and *Act5C* are not affected by *SETDB1* or *wde* GKD.

**Supplementary Figure 3:**
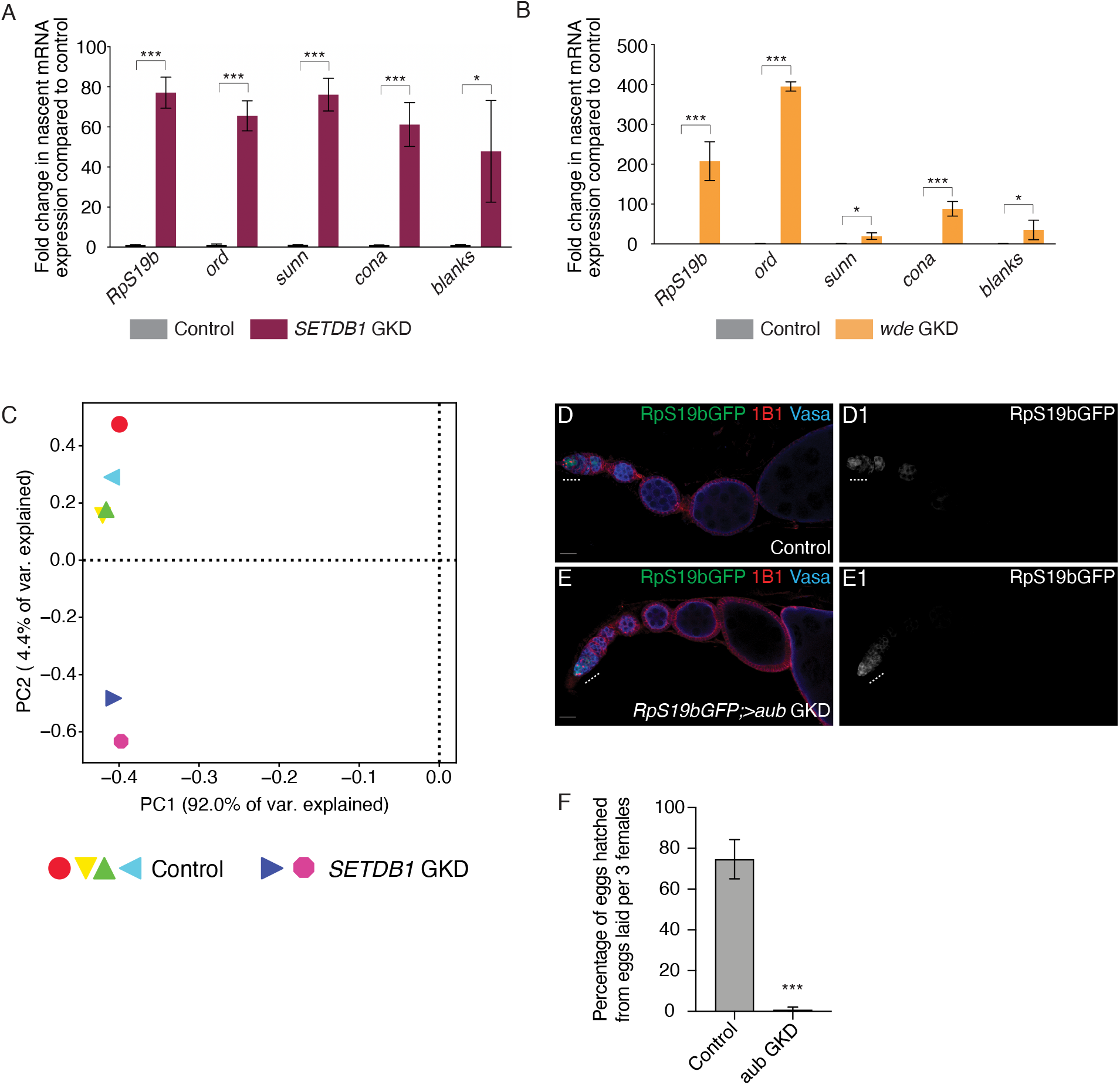
SETDB1/Wde transcriptionally silences expression of a subset of early oogenesis genes. (A) qRT-PCR assaying the pre-mRNA levels of *SETDB1*-regulated target genes, including *RpS19b*, *ord*, *sunn*, *cona* and *blanks* in control and GKD of *SETDB1* shows that these genes are upregulated (n=3, *** indicates p<0.001, Error bars are SEM, Student’s t-Test). (B) qRT-PCR assaying the pre-mRNA levels of *Wde*-regulated target genes, including *RpS19b*, *ord*, *sunn*, *cona* and *blanks* in control and GKD of *Wde* shows that pre-mRNA levels of these genes are upregulated (n=3, * = p ≤ 0.05, ** = p<0.01, *** = p<0.001, Error bars are SEM, Student’s t-Test). (C) Principal Component Analysis (PCA) comparing CUT&RUN data sets for control and *SETDB1* GKD. (D-E1) Ovariole from control *RpS19b-*GFP (D-D1) and GKD of *aub* (E-E1) stained for GFP (green, right grayscale), Vasa (blue) and 1B1 (red). Depletion of this gene shows normal development of the egg chambers and there was no ectopic expression of *RpS19b-*GFP in the egg chambers suggesting SETDB1-mediated silencing of *RpS19b-GFP* is independent of piRNA pathway. (F) Fertility assay of *aub* GKD indicating there was significant decrease in number of adult flies that eclosed from the eggs laid by *aub* GKD flies compared to those from control flies (n=3 trials). *** = p < 0.001, Tukey’s post-hoc test after one-way ANOVA.

**Supplementary Figure 4:**
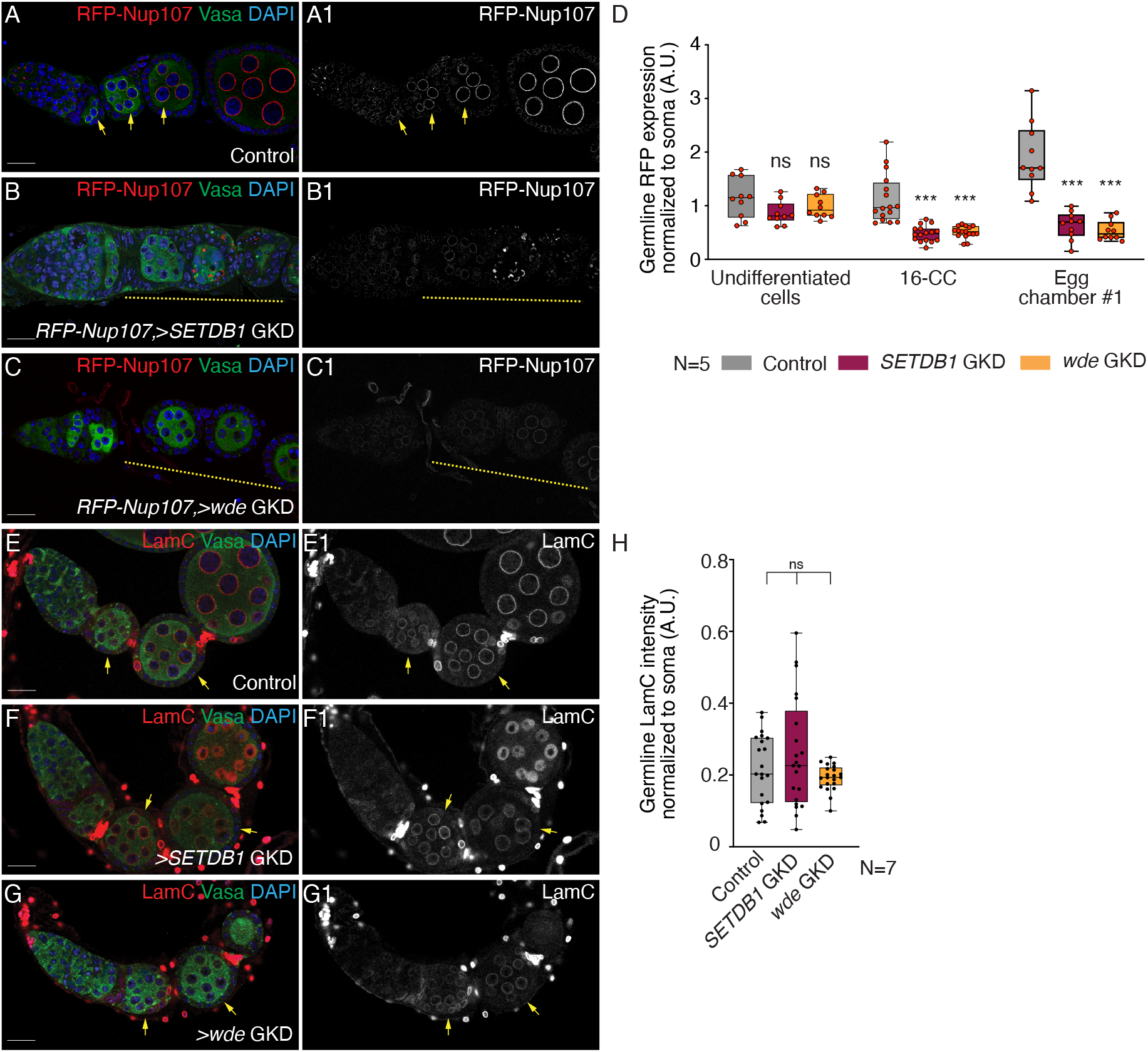
SETDB1/Wde promotes NPC formation without affecting Lamin C. (A-C1) Ovariole of control *RFP-Nup107* A-A1), GKD of *SETDB1* (B-B1) and *wde* (C-C1) stained for RFP (red, right grayscale), Vasa (green) and DAPI (blue). Depletion of *SETDB1* or *wde* shows lower expression of RFP in the egg chambers (yellow line) suggesting *SETDB1*/*wde* regulates expression of Nup107. (D) A.U. quantification of RFP level in the germline normalized to soma in *SETDB1-* and *wde*-GKD ovaries compared to control. Statistical analysis was performed with Dunnett’s multiple comparisons test; N= 5 ovariole; ns = p>0.05, * = p ≤ 0.05, ** = p<0.01, *** = p<0.001. (E-G1) Ovariole of control *UAS-Dcr2;NG4NGT* (E-E2), GKD of *SETDB1* (F-F2) and *wde* (G-G2) stained for LamC (red, right grayscale), Vasa (green) and DAPI (blue). Depletion of *SETDB1* or *wde* shows similar expression of LamC in the egg chambers suggesting *SETDB1* or *wde* depletion does not affect expression of LamC. (H) A.U. quantification of LamC level in the germline normalized to soma in *SETDB1-* and *wde*-GKD ovaries compared to *UAS-Dcr2;NG4NGT* control. Statistical analysis was performed with Dunnett’s multiple comparisons test; N= 7 ovariole; ns = p>0.05.

**Supplementary Figure 5:**
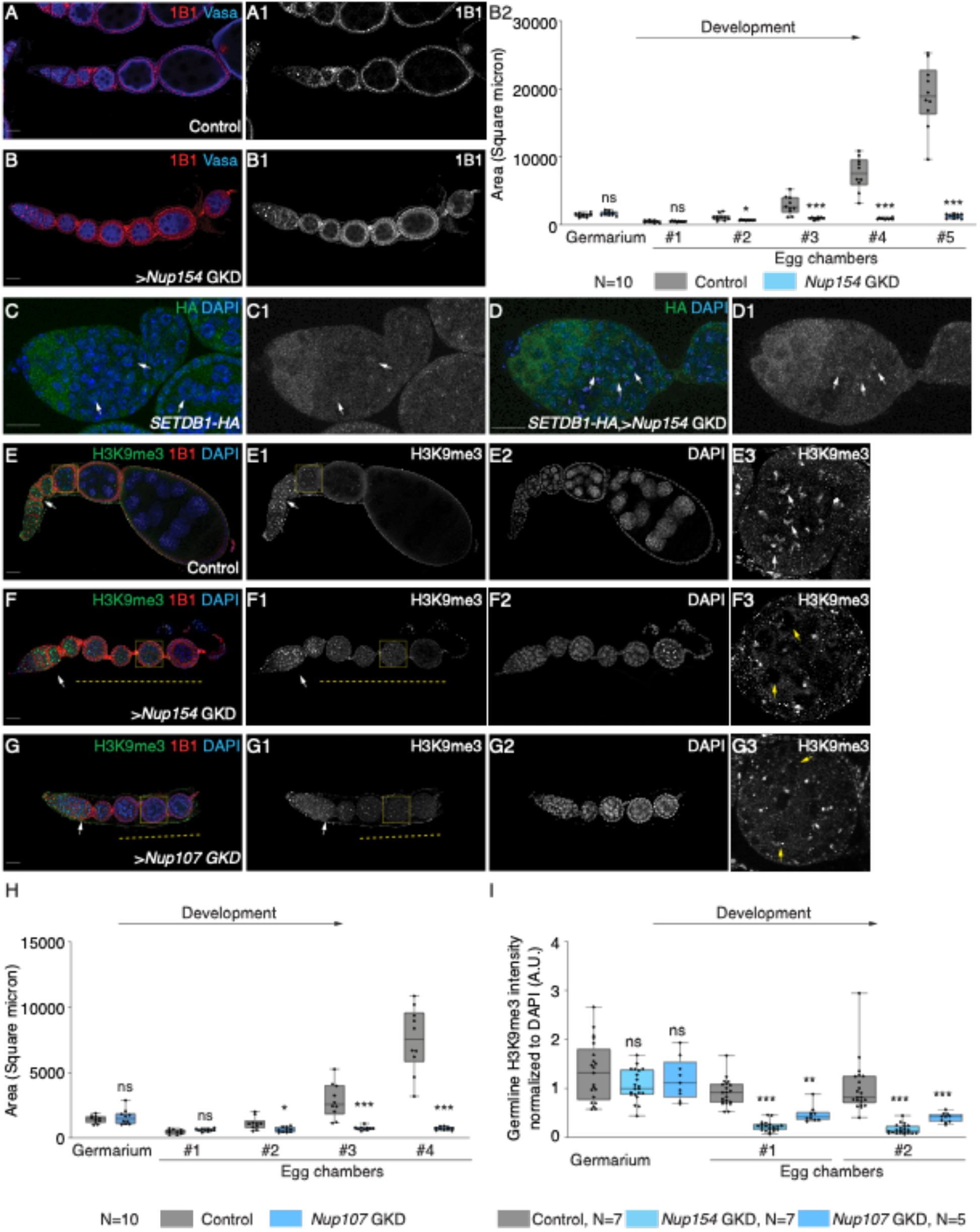
NPC is required for maintaining heterochromatin. (A-B2) Ovariole of control *UAS-Dcr2;NG4NGT* (A-A1) and GKD of *Nup154* (B-B1) stained for 1B1 (red, right grayscale) and Vasa (blue). Control shows normal development of egg chambers while *Nup154* GKD shows egg chambers that do not grow. (B2) Quantification of area of germarium and egg chambers during development in ovaries depleted of *Nup154* (blue) compared to control ovaries (gray). Statistical analysis was performed with Student’s t-test; N= 10 ovarioles; ns = p>0.05, * = p ≤ 0.05, ** = p<0.01, *** = p<0.001. (C-D) Germaria of flies carrying HA tagged SETDB1 (C-C1) and GKD of *Nup154* (D-D1) stained for HA (green, right grayscale) and Vasa (blue). White arrows point at nuclear HA. Depletion of germline *Nup154* shows HA is present in the nucleus. (E-G3) Ovariole and egg chamber of control *UAS-Dcr2;NG4NGT* (E-E3), GKD of *Nup154* (F-F3) and *Nup107* (G-G3) stained for H3K9me3 (green, right grayscale), DAPI (blue, right grayscale) and 1B1 (red). Control shows H3K9me3 expression throughout oogenesis in the germline. Depletion *Nup154* and *Nup107* results in decreased H3K9me3 (yellow dotted line) after differentiation in the egg chambers. Late-stage egg chamber (yellow dotted squares) images show decreased or loss of H3K9me3 in *Nup154* and *Nup107* GKD nurse cells (yellow arrows). (H) Quantification of area of germarium and egg chambers during development in ovaries depleted of *Nup107* (blue) compared to control ovaries (gray). Statistical analysis was performed with Student’s t-test; N= 10 ovarioles; ns = p>0.05, * = p ≤ 0.05, ** = p<0.01, *** = p<0.001. (I) Quantification of H3K9me3 levels in the germline normalized to DAPI level in ovaries depleted of *Nup154* (blue) and *Nup107* (blue) compared to control ovaries (gray). Statistical analysis was performed with Dunnett’s multiple comparisons test; ns = p>0.05, *=p<0.05, ** = p<0.01, *** = p<0.001.

**Supplementary Figure 6:**
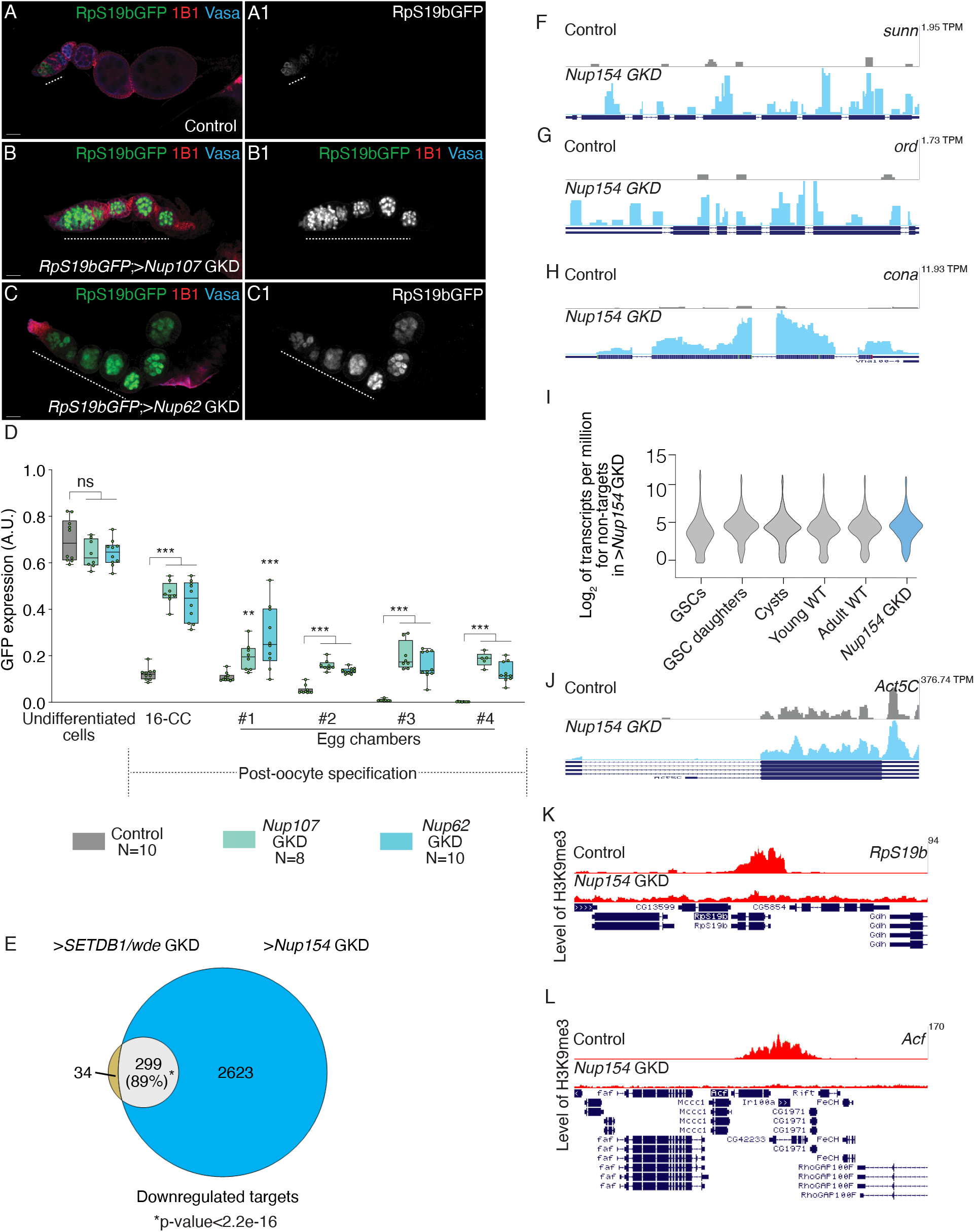
Nup154 represses the early oogenesis genes by promoting their heterochromatinization. (A-C1) Ovariole from control *RpS19b-*GFP (A-A1), GKD of *Nup107* (B-B1) and *Nup62* (C-C1) stained for GFP (green, right grayscale), Vasa (blue) and 1B1 (red). Depletion of these *Nups* shows characteristic phenotype where the egg chambers do not grow and there is ectopic expression of *RpS19b-*GFP in the egg chambers (white dashed line). (D) A.U. quantification of ectopic RpS19b-GFP expression in the germarium and egg chambers with development in ovaries of *Nup107* (teal) and *Nup62* (light blue) GKD compared to control ovaries (gray). Statistical analysis was performed with Dunnett’s multiple comparisons test; ns = p>0.05, ** = p<0.01, *** = p<0.001. (E) Venn diagram of down regulated overlapping genes from RNA-seq of *SETDB1* and *wde* regulated genes with *Nup154* GKD ovaries compared to *UAS-Dcr2;NG4NGT*. 299 down regulated targets are shared between *SETDB1, wde* and *Nup154* GKD, suggesting that Nup154 and SETDB1 function in co-regulating a specific set of genes. (F-H) RNA-seq track showing that synaptonemal complex members *sunn, ord* and *cona* are upregulated upon germline depletion of *Nup154.* (I) Violin plot of mRNA levels of the genes not regulated by *Nup154* in ovaries enriched for GSCs, cystoblasts, cysts, and young and adult whole ovaries, showing that the non-targets of *Nup154* are not silenced in the ovaries compared to cyst stages and whole ovaries. Statistical analysis performed with Hypergeometric test; *** indicates p<0.001. (J) RNA-seq track showing that upon germline depletion of *Nup154, Act5C* is unaffected. (K-L) Tracks showing level of H3K9me3 on target genes. H3K9me3 is depleted on *Nup154* targets *RpS19b* and *Acf* respectively.

